# Phosphorylation of the C-terminus of PI4KA inhibits lipid kinase activity

**DOI:** 10.64898/2026.03.06.710086

**Authors:** Alexandria L Shaw, Sophia Doerr, Hunter G Nyvall, Meredith L Jenkins, Sushant Suresh, Calvin K Yip, Scott D Hansen, John E Burke

## Abstract

Phosphatidylinositol 4-kinase alpha (PI4KA) is a lipid kinase that generates phosphatidylinositol 4-phosphate (PI4P) from phosphatidylinositol (PI) at the plasma membrane (PM). PI4P generated by PI4KA is essential for both plasma membrane identity and for PIP_2_ and PIP_3_ signalling driven by the PLC and PI3K family of enzymes. While the structure of PI4KA is known, the regulatory mechanisms that control its activity are undefined. Here, we discovered PI4KA can be inhibited through phosphorylation of Y2090 in the kinase domain, which can be phosphorylated by multiple tyrosine kinases. Y2090 phosphorylation causes local conformational changes in the kα12 C-terminal helix. PI4KA activity is predominantly controlled through membrane recruitment by EFR3, with Y2090 phosphorylation inhibiting EFR3 bound PI4KA. Phosphorylation of the kα12 C-terminal helix is found in PI3Ks and PI4Ks, suggesting an evolutionarily conserved regulatory mechanism for diverse phosphoinositide kinases. Overall, our work reveals novel molecular insight into inhibitory post-translational regulation of PI4KA.

## Introduction

Type III phosphatidylinositol 4-kinase α (further referred to by its gene name *PI4KA*) is an essential lipid kinase that facilitates the transfer of phosphate onto the 4 position of phosphatidylinositol (PI) generating phosphatidylinositol 4-phosphate (PI4P)^1^. PI4KA is a master regulator of plasma membrane (PM) identity through the generation of plasma membrane (PM) asymmetry through exchange of PM PI4P for ER phosphatidylserine by the ORP5/8 lipid transfer proteins^2^. The generation of PI4P is also required for PI(4,5)P_2_ and PI(3,4,5)P_3_ synthesis, which are important lipid second messengers involved in phospholipase C (PLC) (calcium signalling) and PI3K/Akt (cellular growth, proliferation, survival, and metabolism) signalling pathways^3^. PI4KA consists of multiple distinct domains. At the N-terminus is an α-solenoid (commonly referred to as the horn) followed by a domain that facilitates dimerization of two PI4KA proteins (called the dimerization domain). Similar to class I PI3Ks, PI4KA has a C-terminal bi-lobal kinase domain that is cradled by its helical domain. Functional PI4KA primarily exists as a heterotrimeric complex with the regulatory proteins TTC7 and FAM126 (both of which have A and B isoforms). Two PI4KA/TTC7/FAM126 heterotrimers associate to form a homodimer^4–6^.

A defining feature of phosphoinositide kinases (class I-III PI 3-kinases (PI3Ks) and PI4KA/B) regulation are their recruitment to membrane surfaces through dedicated lipid and protein binding modules. An established regulator of PI4KA membrane recruitment is the palmitoylated EFR3(A/B) protein^7^. It anchors PI4KA in proximity to its lipid substrate through contacts between EFR3’s C-terminus and both regulatory proteins TTC7 and FAM126^8–10^. Previous work has suggested an alternative assembly of EFR3 with transmembrane protein TMEM150 to localise the PI4KA complex to the plasma membrane in the absence of TTC7/FAM126^11^, however, the molecular basis for complex assembly is unknown. Calcium signalling driven by PLC mediated hydrolysis of PI(4,5)P_2_ has also been linked to PI4KA regulation, with the palmitoylated CNAβ1 calcineurin isoform making direct interactions with PI4KA and FAM126A. Functionally, it has been demonstrated that following depletion of the PI(4,5)P_2_ pool through PLC signalling, CNAβ1 regulates PI4KA activity by depletion of PI4KA complex phosphorylation^12,13^. Regulation of PI4KA activity could play an important role in shaping plasma membrane signalling dynamics, as inhibition of PI4KA disrupts the oscillatory behaviour of PI4P, PI(4,5)P_2_, and PI(4,5)P_2_ dependent effectors at the plasma membrane^14^. PI4KA has been found to be phosphorylated in many phosphoproteomic MS experiments^15^, however, the effect of these phosphosites on PI4KA activity has not been characterised.

PI4KA shares an evolutionary lineage with PI4KB and the class I, II, and III phosphoinositide 3-kinases^16^. Cryo EM analysis of PI4KA has revealed the architecture of how PI4KA assembles with FAM126A and TT7B^4^, as well as how it can associate with its partners EFR3A^9^, and the protein phosphatase calcineurin^13^. In all of these structures the kinase domain of PI4KA adopts a fold similar to PI4KB and the class I, II, and III phosphoinositide 3-kinases. Conserved in all of these enzymes is a predicted kα12 C-terminal helix in the kinase domain that is absolutely required for membrane association and lipid kinase activity^17–19^. In all structures of PI4KA this helix is disordered, with it also being disordered in most PI3K/PI4KB structures, however, recent structural work of p110α bound to membranes shows ordering of this helix and its interaction with membranes^20^. Ordering of this helix upon membrane binding has been proposed for all PI3Ks and PI4Ks, with this being consistent with HDX-MS analysis of class I PI3Ks^19^. This C-terminal helix is amphipathic, with a hydrophobic face that interacts with the membrane core, and a hydrophilic face pointing towards solution. Phosphorylation of residues in the kα12 helix have been found in the class I PI3Ks for p110α, p110β, and p110δ, with all leading to decreased lipid kinase activity^21–23^. This is putatively driven by autophosphorylation in p110β, and p110δ, while p110α is phosphorylated downstream of epinephrine activation of hippo kinase signalling^23^. Overall, this suggests a potential role of kinase domain phosphorylation in regulating PI4KA activity.

The PI4KA-TTC7-FAM126-EFR3 signalling axis is dysregulated in myriad human diseases, with both the upregulation and downregulation of this pathway being implicated^24^. Historically, due to its requirement for hepatitis C viral replication, where its activity is hijacked to produce increased amounts of PI4P at viral replication organelles^25^, extensive efforts were dedicated to develop highly selective ATP-competitive inhibitors. However, the progression of these into the clinic was hindered by the fact that PI4KA is essential to cellular function, with both genetic and pharmacological inhibition of PI4KA being lethal^26^. While it is suggested that PI4KA inhibition may have a role in combination therapies, particularly in Ras driven cancers, reducing toxicity will be challenging^27,28^. Identification of alternative therapeutic avenues for PI4KA related disorders is limited, therefore understanding the molecular basis for regulation by regulatory proteins and post translational modifications is critically important.

To explore how PI4KA is regulated by phosphorylation we have used a multi-pronged approach to identify kinases that directly phosphorylate PI4KA and functionally characterise the effect of these phosphosites. We identified Y2090 and Y1154 as sites that can be modified by multiple tyrosine kinases. Functional assays demonstrated that phosphorylation of Y2090 suppresses lipid kinase activity even when PI4KA is tethered to its membrane substrate by its membrane resident regulatory protein EFR3, whereas Y1154 had no effect on kinase activity. Cryo-EM and HDX-MS revealed the molecular basis for this regulation: pY2090 alters the conformation of a C-terminal helix (kα12) of the kinase domain that plays a critical role in lipid kinase activity. Overall, we have identified a PI4KA inhibitory mechanism driven by phosphorylation of Y2090, which suggests a previously unknown negative feedback input into PI4KA signalling.

## Results

The discovery of PI4KA regulation by the protein phosphatase calcineurin motivated us to analyse previously identified phosphorylation sites in PI4KA and its accessory proteins using PhosphoSitePlus^15^ (**Fig. S1)**. The most frequently reported site was Y1154 in PI4KA (>250 observations), positioned within the dimerization domain at the PI4KA dimer interface helix (**Fig. 1A/E)**. Recent advances in the prediction of tyrosine kinase substrate selectivity^29^ identified lymphocyte-specific protein tyrosine kinase (Lck) as the most efficient kinase for this site. Intriguingly, an interaction of PI4KA with Lck has been previously described, suggesting potential biological relevance of this kinase in PI4KA regulation^30,31^.

**Figure 1.**
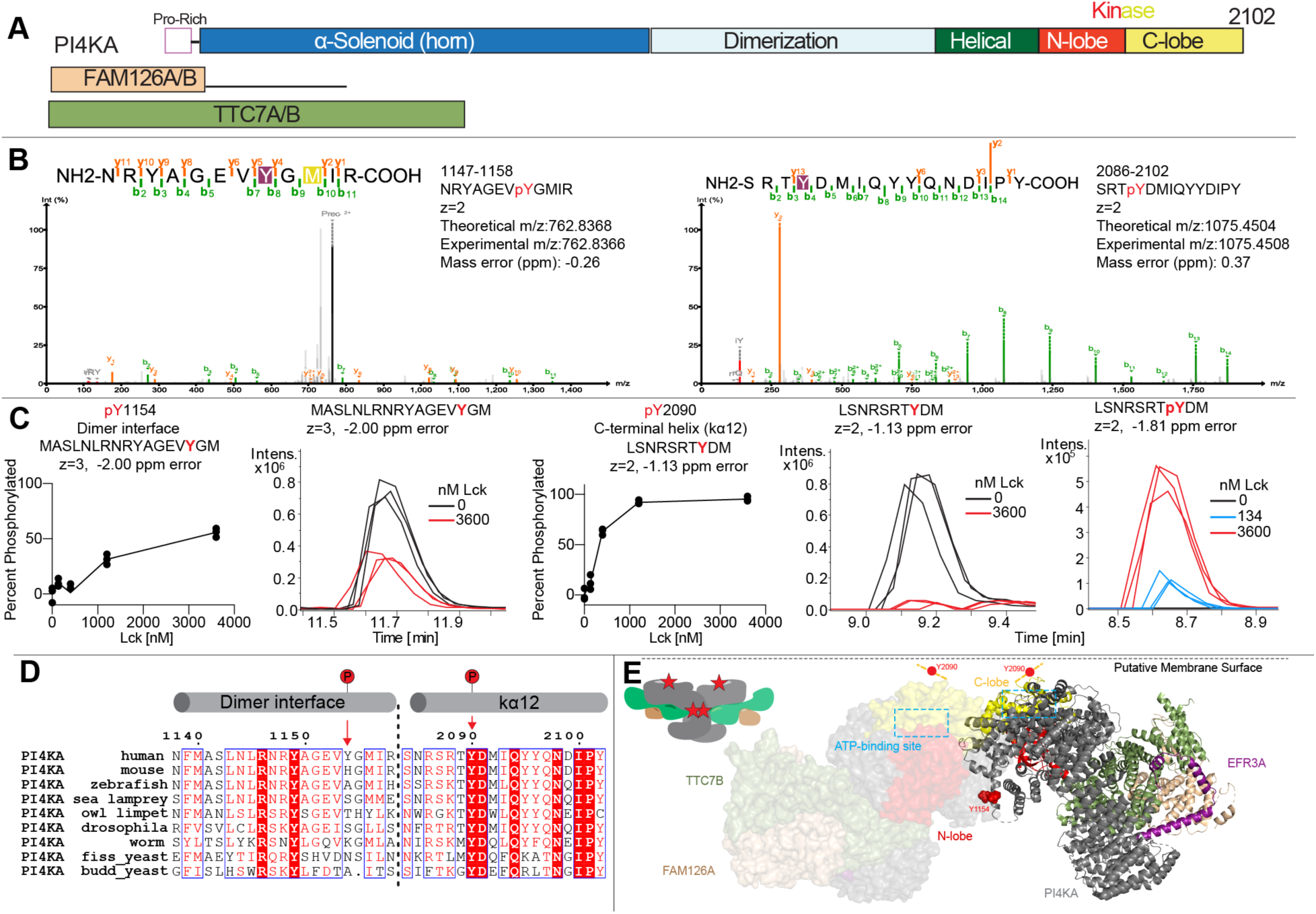
PI4KA can become phosphorylated at tyrosine 1154 and 2090. A. Domain schematics of the lipid kinase PI4KA, highlighting key functional regions and regulatory proteins TTC7 and FAM126A. B. Representative tandem mass spectrometry (MS/MS) spectra of phosphorylated peptides containing pY1154 and pY2090. Peptides were generated by tryptic digestion and analyzed by higher-energy collisional dissociation (HCD) MS/MS. The b- and y-fragments are annotated on the sequence and highlighted in the spectrum. Theoretical and experimentally observed precursor m/z values for each peptide are shown, with ppm error included, additional fragments are included in Fig. S2. C. Phosphorylation quantification of pY1154 and pY2090 levels in _WT_PI4KA upon treatment with varying concentrations of Lck. The percent phosphorylation was quantified by taking the inverse of the loss of non-phosphorylated peptide intensity upon increasing amounts of Lck 225-509 (Data are shown as individual values, n=3 for each peptide). Extracted ion chromatograms of peptides used to quantify phosphorylation of pY1154 and pY2090 are shown in the presence (3.6 µM) and absence of Lck. The intensity of the phosphorylated peptides as well as the full set of extracted ion chromatograms for all Lck concentrations are shown in Fig S3. D. Multiple sequence alignment of PI4KA from *Homo sapiens*, *Mus musculus*, *Danio rerio*, *Petromyzon marinus*, *Lottia gigantea*, *Drosophila melanogaster*, *Caenorhabditis elegans*, *Schizosaccharomyces pombe*, and *Saccharomyces cerevisiae*, showing conservation of the PI4KA Y2090 site and lack of conservation of Y1154 with predicted secondary structure indicated above (phosphorylation sites are indicated by red arrows). E. Structure of PI4KA complex bound to EFR3 (9BAX)_9_ annotated with the identified phosphorylation sites Y1154 and Y2090. The Y2090 site is in a disordered segment located at the C-terminus of PI4KA. The ATP binding site is indicated in a blue box on both PI4KA protomers.

To determine if PI4KA could be directly phosphorylated by Lck, we purified an active construct of Lck (residues 225–509) and co-incubated heterotrimeric PI4KA-TTC7B-FAM126A (hereafter referred to as ^WT^PI4KA), Mg-ATP and a saturating amount of Lck kinase. We then analyzed these samples using LC-MS/MS to unambiguously identify any Lck mediated phosphosites. We identified two phosphorylated sites (Y1154 and Y2090) **(Fig. 1B/S2 and Source Data)** in PI4KA that were unique to the Lck-treated sample, with no identified tyrosine-phosphorylated sites in TTC7B and FAM126A. Next, we treated ^WT^PI4KA with varying concentrations of Lck together *in vitro* to quantify the phosphorylation efficiency of these sites **(Fig. 1C)**. Phosphorylation was quantified from the depletion of the corresponding non-phosphorylated peptide relative to samples lacking Lck. We quantified the intensity of the non-phosphorylated and phosphorylated peptides for pY1154 (**Fig. S3A+B**) and pY2090 (**Fig. S3C+D**) as well as control peptides that eluted at similar times (**Fig. S3E**). Quantification of Y1154 phosphorylation at different concentrations of Lck showed a plateau at ∼50% **(Fig. 1C)**, suggesting that only one protomer within the dimer was modified. Intriguingly, Lck treatment robustly phosphorylated Y2090, which is located in the disordered terminus of the kinase domain, which forms the putative membrane binding kα12 helix **(Fig. 1C)**. Phosphorylation of this site was more efficient, plateauing at ∼90%. This site has been detected in previous phosphoproteomic analysis of PI4KA^15,32–36^, but not at the same frequency as Y1154. Evolutionary analysis of these sites revealed that Y1154 is weakly conserved, while the Y2090 in the C-terminus is strictly conserved from *<u>Homo sapiens</u>* <u>to yeast</u> **(Fig.1D)**.

To further understand the specificity of Y1154 and Y2090 phosphorylation by tyrosine kinases, we utilized the kinase prediction tool from the kinase library (https://kinase-library.mit.edu/home) to predict the kinases with the highest specificity for the Y1154 and Y2090 sites **(Fig. 2A)** ^29,37^. Y1154 was strongly ranked for recognition by Src-family kinases, with Lck, HCK, and SRC among the top candidates. These kinases are central regulators of signaling: Lck, which is essential for T-cell signalling ^38^, HCK which regulates cell adhesion, proliferation, migration, invasion, apoptosis, and angiogenesis in hematopoietic stem cells ^39^, and SRC which is a proto-oncogene that regulates cellular survival, proliferation, angiogenesis, and motility ^40,41^. In contrast, Y2090 was predicted to be targeted by a distinct set of kinases, with DDR1, DDR2 (DDR family), and Anaplastic lymphoma kinase (ALK) (LTKR family) among the highest-ranked. DDR1/2 are non-integrin collagen receptors essential for developmental processes ^42,43^, while ALK is an oncogenic receptor tyrosine kinase expressed in the nervous system ^44^. Notably, Src-family members were ranked much lower for Y2090 (Lck, 36th; Src, 47th out of 78), highlighting the divergent kinase specificity of the two sites. Interestingly, the insulin receptor, which is a major activator of the PI3K-AKT signalling cascade, was also predicted, ranking 9th for Y2090 and 25th for Y1154. To assess kinase specificity toward PI4KA, we treated the ^WT^PI4KA with a fixed amount (1 µg) of Lck, SRC, or INSR (SDS-PAGE gel in Source Data) and quantified phosphorylation at Y1154 and Y2090 by LC-MS **(Fig. 2B)**. Lck phosphorylated Y1154 to ∼30% and Y2090 to ∼90%, with Lck showing the largest difference in phosphorylation levels between the two sites **(Fig. 2C)**. SRC phosphorylated Y1154 to ∼15% and ∼40% at Y2090 **(Fig. 2D)** where treatment with INSR kinase Y2090 reached ∼30% with no observed phosphorylation of Y1154 **(Fig. 2E)**. Additionally, across all three kinases, Y2090 consistently exhibited higher phosphorylation than Y1154, suggesting that Y1154 is comparatively less accessible to modification **(Fig. 2C-D)**. This is consistent with the fact that the C-terminus is disordered, and it likely more available for phosphorylation than Y1154 which is in an ordered region of the protein.

**Figure 2.**
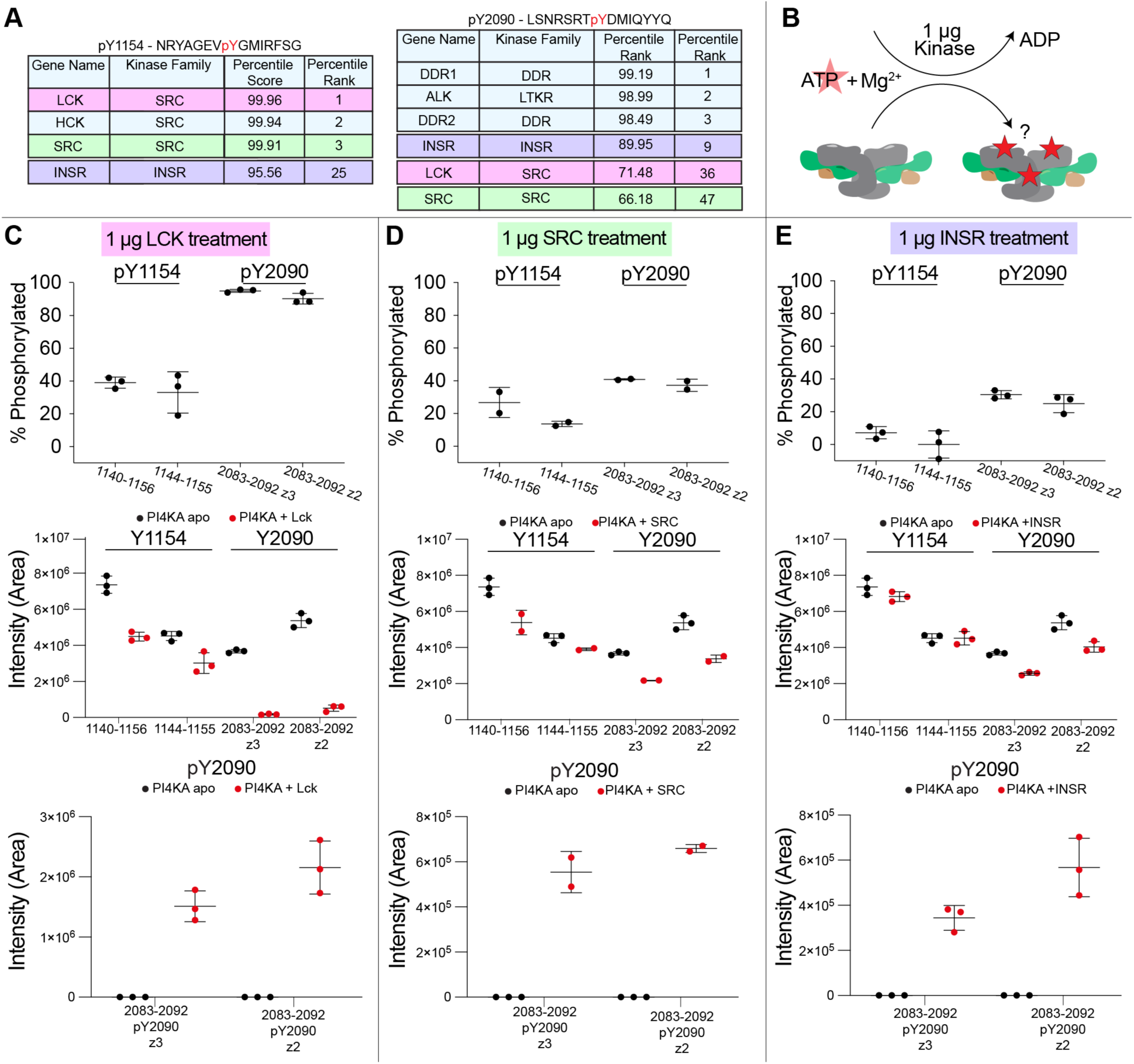
Direct phosphorylation of the PI4KA complex by tyrosine kinases. A. Top predicted kinases, using The Kinase Library to phosphorylate Y1154 and Y2090 in PI4KA. B. Diagram of reaction setup. PI4KA was mixed with ATP-MgCl_2_ mixture and added to 1 ug of different tyrosine kinases. Phosphorylation was quantified using LC-MS. C/D/E. (Top)Phosphorylation quantification of _WT_PI4KA (2 µM) Y1154 and Y2090 upon treatment of ^WT^PI4KA complex with ATP/MgCl_2_ and 1 µg (C) Lck (225-509) (1.5 µM), (D) SRC (FL) (2.37 µM) (3.02 µM), and (E) INSR (KD) compared to untreated. Phosphorylation was quantified by taking the inverse of the loss of unphosphorylated peptide intensity across peptide(s). (Middle) Intensity (Area extrapolated from extract ion chromatograms) of two unique PI4KA peptides spanning Y1154 and one peptide with two charge states spanning Y2090 comparing PI4KA apo to (C) Lck, (D) SRC, and (E) INSR treatment. (Bottom) Increase in pY2090 containing peptides (1 peptide with two charge states) upon treatment with (C) Lck, (D) SRC, and (E) INSR. Additional information regarding the peptides tracked in this figure can be found in the source data. (n=3 for apo and INSR and Lck treatments, n=2 for SRC treatment, independent replicates, error bars represent SD).

### Assessing the effect of phosphorylation on PI4KA activity

To biochemically characterize the role of tyrosine phosphorylation in PI4KA regulation, we purified phosphorylated PI4KA through *in vitro* Lck phosphorylation (Phosphorylated and non-phosphorylated PI4KA/TTC7B/FAM126A 1-308 complexes will be referred to as ^WT^PI4KA^pY1154,pY2090^ or ^WT^PI4KA, respectively). ^WT^PI4KA was treated with active Lck (225-509) in the presence of ATP and MgCl_2_, followed by size exclusion chromatography (SEC) to remove residual Lck. Both phosphorylated and non-phosphorylated PI4KA complexes eluted from gel filtration at a volume consistent with a dimeric complex (∼730 kDa) with similar purity assessed by SDS-PAGE **(Fig. S4)**. With these elution profiles suggesting that phosphorylation does not affect PI4KA’s ability to dimerize. Phosphorylation of purified PI4KA complex was quantified by LC-MS comparing the intensity of the non-phosphorylated peptide in the non-phosphorylated sample to the phosphorylated sample, with Y1154 being ∼50% phosphorylated and Y2090 being ∼90% phosphorylated **(Source Data)**.

To evaluate how phosphorylation alters PI4KA’s lipid kinase activity, we conducted kinase assays using 100% PI vesicles as substrate and measured ATP turnover. ^WT^PI4KA^pY1154,pY2090^ showed a ∼15-fold decrease in PI4KA activity compared to unphosphorylated **(Fig. 3A/B)**. Importantly, the intrinsic ability for PI4KA to turn over ATP in the absence of lipid substrate, termed ATPase activity, is unaffected by phosphorylation of Y1154 and Y2090, suggesting phosphorylation does not decrease the intrinsic ATP turnover of PI4KA, but instead likely alters either lipid engagement or catalysis **(Fig. 3B)**. It is important to note that even with extensive optimisation we could never generate a 100% phosphorylated version of PI4KA, so the 15-fold decrease may be an underestimation of the effect of phosphorylation due to the ∼10% of unphosphorylated complex present in the sample. Although phosphorylation clearly had a substantial effect on kinase activity, it remained unclear at this stage whether the change in lipid kinase activity was attributable to phosphorylation of Y1154, Y2090, or both residues.

**Figure 3.**
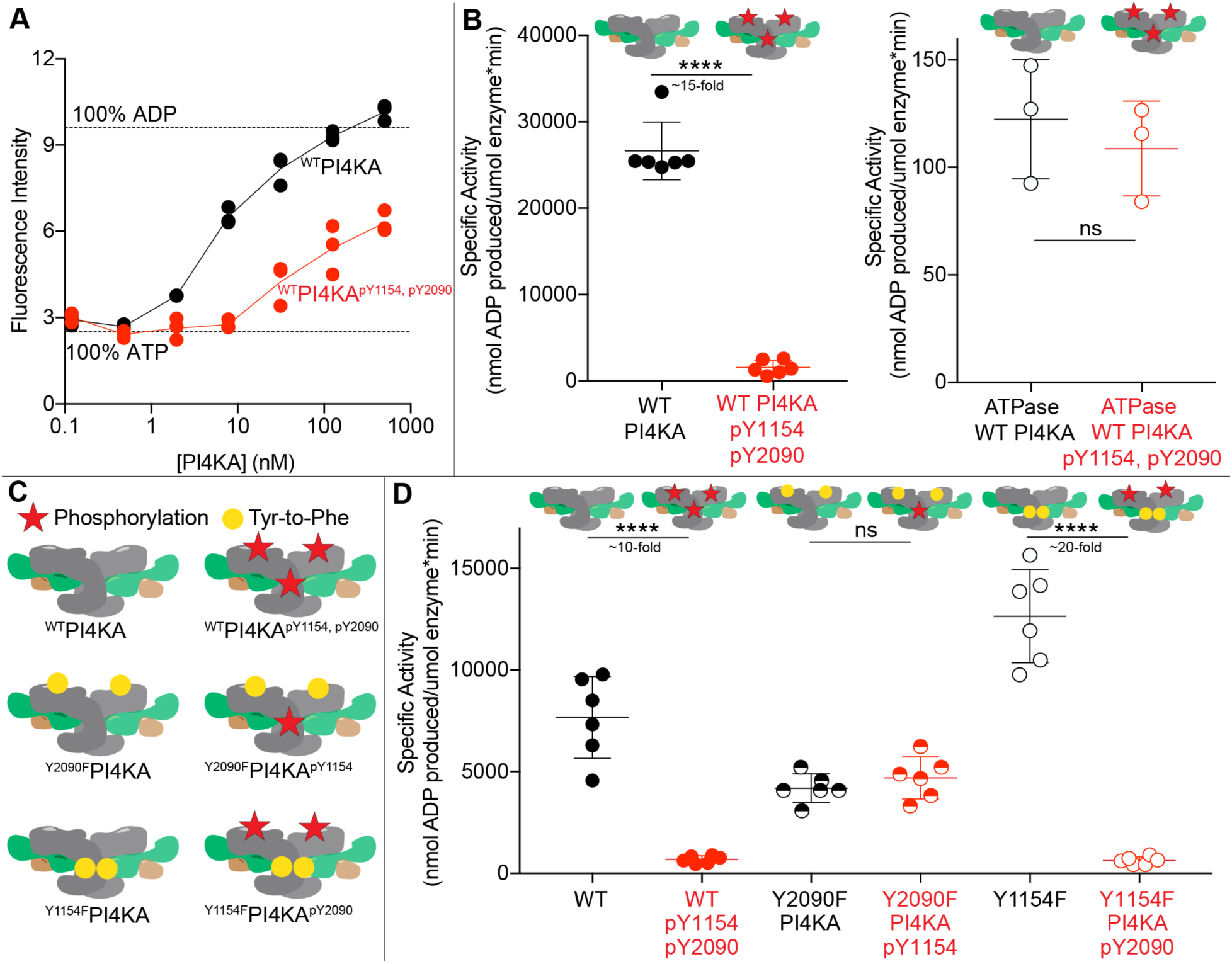
Phosphorylation of Y2090 drives a large decrease in PI4KA lipid kinase activity. A. Measurement of ATP turnover of ^WT^PI4KA (black) and ^WT^PI4KA^pY1154,pY2090^ (red) in the presence of 100% PI vesicles. Experiments were performed with 0.12-500 nM and a final vesicle concentration of 0.5 mg/ml. (Independent replicates; data are presented as mean values, and error bars are SD, n=3) B. Specific activity values for ^WT^PI4KA and ^WT^PI4KA^pY1154,pY2090^ (L) with 100% PI vesicles and (R) in the absence of vesicles, called ATPase activity. Specific activity was calculated using the concentrations where kinase activity was in range (1.95 and 7.81 nM for ^WT^PI4KA with 100% PI vesicles, 31.25 and 125 nM for ^WT^PI4KA^pY1154,pY2090^ with 100% PI vesicles, and 500 nM for ATPase). (Independent replicates; data are presented as individual points, and error bars are SD, n=3 for each concentration). Two-tailed *t* test P values are represented by; ns and **** <0.0001. C. Cartoon schematics of phosphorylated PI4KA constructs used in panel D. Red star denotes phosphorylation (pY1154 ∼40-50%, pY2090 ∼90%). Yellow circle denotes Tyr-to-Phe mutation at Y2090 or Y1154. D. Specific activity values for ^WT^PI4KA and ^WT^PI4KA^pY1154,pY2090^, ^Y2090F^PI4KA, ^Y2090F^PI4KA^pY1154^, ^Y1154F^PI4KA, and ^Y1154F^PI4KA^pY2090^ with 100% PI vesicles. Specific activity was calculated using the concentrations where kinase activity was in range (7 and 17.5 nM for ^WT^PI4KA, ^Y2090F^PI4KA, ^Y2090F^PI4KA^pY1154^ and ^Y1154F^PI4KA and 125 and 312.5 nM for ^WT^PI4KA^pY1154,pY2090^ and ^Y1154F^PI4KA^pY2090^. (Technical replicates; individual data points are shown, and error bars are SD, n=3 for each concentration). Two-tailed *t* test P values are represented by; ns and **** <0.0001.

Determining the molecular mechanism governing this significant loss-of-function in PI4KA required generating PI4KA complexes that were monophosphorylated (either pY1154 or pY2090). We thus generated tyr-to-phe mutations for both Y1154 (Y1154F) and Y2090 (Y2090F) and treated these with Lck **(Fig. 3C)**. Once again these were co-expressed with TTC7B and FAM126A 1-308 in *Sf9* cells and purified in the same manner as ^WT^PI4KA and ^WT^PI4KA^pY1154,pY2090^ complexes. All complexes eluted from size exclusion chromatography columns at a volume consistent with the dimeric heterotrimer complex and had comparable purity determined by SDS-PAGE (**Fig.S4**). The Y2090F construct had ∼40% phosphorylation of pY1154, and the Y1154F construct had ∼95% phosphorylation of Y2090 **(Source data)**. The only difference in observed phosphorylation compared to WT complexes was that we observed ∼20% phosphorylation in a peptide containing Y2096 and Y2102 in the Y2090F construct **(Source data)**. These proteins will be referred to as ^Y2090F^PI4KA, ^Y2090F^PI4KA^pY1154^, ^Y1154F^PI4KA, and ^Y1154F^PI4KA^pY2090^, respectively.

We carried out kinase assays using 100% PI substrate for all PI4KA complexes. This revealed no significant difference in lipid kinase activity between ^Y2090F^PI4KA and ^Y2090F^PI4KA^pY1154^ **(Fig. 3D)**, indicating that the dimer interface PTM site does not alter lipid kinase activity. In contrast, kinase assays comparing ^Y1154F^PI4KA and ^Y1154F^PI4KA^pY2090^ showed a ∼20-fold decrease in PI4KA activity, which was similar to that observed for ^WT^PI4KA^pY1154,pY2090^ **(Fig. 3D)**. Therefore, phosphorylation of the C-terminus of PI4KA at Y2090 is responsible for the observed decrease in PI4KA kinase activity.

### Cryo-EM and HDX-MS analysis of the PI4KA complex

To understand how phosphorylation at Y2090 and Y1154 influence PI4KA complex dynamics, we employed cryogenic electron microscopy (cryo-EM), hydrogen-deuterium exchange mass spectrometry, and AlphaFold3 modelling.

We conducted cryo-EM analysis of ^WT^PI4KA^pY1154,pY2090^. Prior to vitrification, ^WT^PI4KA specimen was treated with Lck (225-509) and ATP/MgCl_2_. Protein phosphorylation was followed by a dose of BS^3^ crosslinker to stabilize the complex immediately prior to vitrification^9,10,13^. LC-MS analysis of this sample prior to the addition of crosslinker was used to quantify phosphorylation of Y1154 and Y2090 to be ∼40% and ∼95% phosphorylated, respectively. Quantification of this sample was conducted by comparing the intensity of the phosphorylated peptide to the overall peptide (phosphorylated and unphosphorylated) intensity, as there was no direct comparison made without Lck (**Source Data**). Single particle analysis resulted in the refinement of 239,639 particles resulting in a 3D reconstruction at a nominal resolution of 2.98 Å **(Fig. 4A/S5)**. Fortunately, the highest local resolution was observed at the dimerization domain interface where pY1154 is located **(Fig. S5)**. However, the interpretation of the EM reconstruction of the dimerization interface is complicated by two main factors. First, residue Y1154 was ∼40% phosphorylated. Second, the presence of C2 symmetry, which can complicate EM processing algorithms resulting in unique density present in one symmetric unit being cancelled out by a symmetric unit lacking this feature. As a result, the EM map likely shows an ensemble of orientations of side chains at this interface at a higher threshold, representing a mixture of unphosphorylated and phosphorylated Y1154.

**Figure 4.**
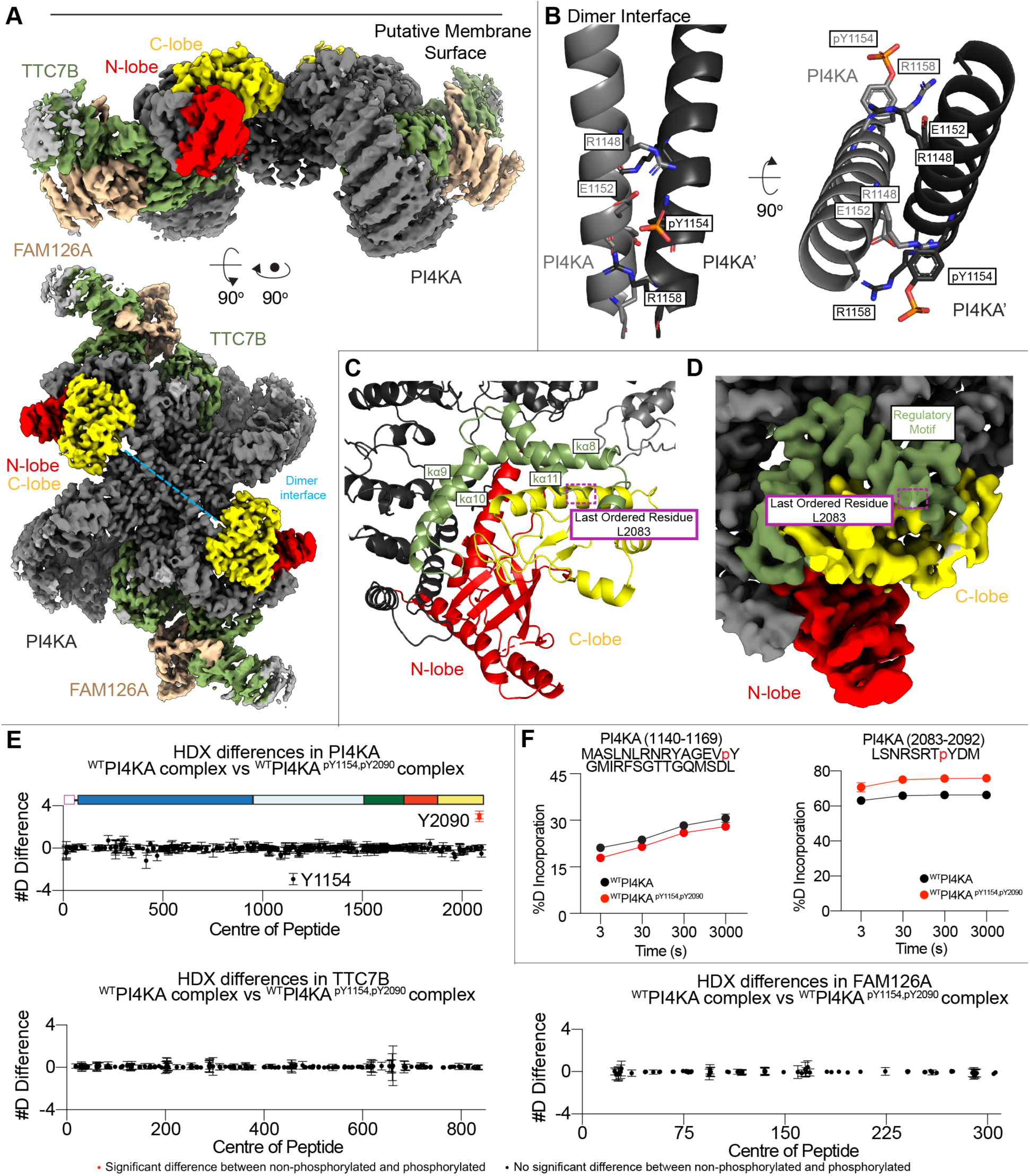
Cryo-EM structure of ^WT^PI4KA^pY1154,pY2090^ complex. A. Cryo-EM density map of ^WT^PI4KA^pY1154,pY2090^. B. Cartoon representation of the dimerization interface, highlighting the coordination of E1152 and R1148 in PI4KA with pY1154 and R1158 in PI4KA’. C. Cryo-EM model of the ^WT^PI4KA^pY1154,pY2090^ kinase domain. The bi-lobal kinase domain is shown in red (N-lobe) and yellow (C-lobe), highlighting the conserved regulatory arch in green. Dotted box represents the last ordered residue in our cryo-EM model, stopping before residues corresponding to the kα12 helix. D. Cryo-EM density map of the kinase domain in the ^WT^PI4KA^pY1154,pY2090^ PI4KA complex coloured according to panel C. E. Sum of the number of deuteron difference of PI4KA, TTC7B, and FAM126A upon phosphorylation of pY1154 and pY2090, analyzed over the entire deuterium exchange time course for the PI4KA complex. Each point is representative of the center residue of an individual peptide. Peptides that met the significance criteria (defined as >5%, 0.4 Da, and p < 0.01 in an unpaired two-tailed t test at any time point) coloured red. Error is shown as the sum of standard deviations across all four timepoints, n=3. F. Select deuterium exchange time courses of PI4KA peptides that showed changes (left-failed to pass all significance criteria, right-significant change) in deuterium exchange when phosphorylated. Error is shown as SD, n=3.

When building the molecular model, additional density was observed adjacent to the hydroxyl of Y1154 in both copies of the PI4KA dimer, allowing for a phosphate group to be modelling on this tyrosine **(Fig. 4B/ S6)**. We compared our model of ^WT^PI4KA^pY1154, pY2090^ to published cryo-EM models of the PI4KA complex (PDB:9BAX, PDB:9B9G). These comparisons yielded overall RMSD values of 0.379 Å and 1.794 Å, respectively. To identify potential perturbations at the dimerization interface, we focused on differences between PDB:9BAX, PDB:9B9G, and ^WT^PI4KA^pY1154,pY2090^ in this region. For residues 1100-1400, RMSD values were 0.267 Å and 0.578 Å relative to PDB:9BAX and PDB:9B9G, respectively. For the isolated dimerization helix (residues 1136-1159), RMSD values were 0.261 Å and 0.411 Å, respectively. Key residues surrounding Y1154 include R1148, E1152, and R1158. Their orientations exhibit a high degree of similarity across the structures, with only a slight shift observed for R1158 (**Fig. S6).** The Y1154 residue of PI4KA is in close proximity to Y1154 from the other protomer, possibly explaining why phosphorylation of the 1^st^ Y1154 prevents phosphorylation of the 2^nd^ Y1154 through changing the local environment and preventing Lck engagement. Overall, these comparisons suggest that phosphorylation of Y1154 has only a minor effect on the conformation of PI4KA.

The PI4KA and PI4KB enzymes along with the class I,II, and III PI3Ks all contain a regulatory motif in their kinase domain that is C-terminal to the activation loop. This region is composed of helices kα7–kα12, with the final helix being disordered in most PI4K and PI3K structures. Located in the kα12 helix of the kinase domain regulatory motif is Y2090. In previously reported PI4KA cryo-EM reconstructions, kα12 was not structurally resolved. Similarly, our cryo-EM map of ^WT^PI4KA^pY1154,pY2090^ lacked density for the kα12 helix (**Fig. 4C)**. AlphaFold3 predicted formation of the kα12 helix in ^WT^PI4KA^pY2090^, however this prediction has low confidence (**Fig. S7)**. The predicted local distance difference test (pLDDT) score for this region was below 60%, and the predicted aligned error (PAE) between the kα12 helix and the rest of the PI4KA complex was high, showing low spatial confidence. (**Fig. S7).**

The lack of density in the cryo-EM reconstruction of the kα12 helix required an alternative method to elucidate the effect of phosphorylation on kα12 dynamics. To address this question, we utilised hydrogen-deuterium exchange mass spectrometry (HDX-MS). HDX-MS reports on changes in protein conformational dynamics by measuring the exchange of backbone amide hydrogens with deuterated solvent, a process primarily governed by secondary structure. Exchange patterns provide insight into local protein dynamics upon different conditions, such as phosphorylation. Deuterium incorporation is localised at the peptide level through proteolytic digestion. HDX-MS experiments comparing ^WT^PI4KA to ^WT^PI4KA^pY1154,pY2090^ were carried out at 1.2 uM (12 pmol), with deuterium incorporation being measured over four exchange time points (3, 30, 300, and 3000 s at 20°C). The full raw deuterium incorporation for the HDX-MS experiment is provided in the source data. Coverage of all proteins was over 88% with all HDX-MS processing statistics available in the **source data**. Overall, we detected only one significant difference in H/D exchange in PI4KA. This occurred in a peptide spanning residues 2083-2092 (PI4KA kα12) upon Y2090 phosphorylation **(Fig. 4E/F)**. However, no large conformational changes distant from the direct phosphorylation site were observed. We also observed a subtle non-significant decrease in deuterium exchange in a peptide spanning residues 1140-1169 (PI4KA dimerization interface, Y1154) when phosphorylated **(Fig. 4E/F)**. It is important to note that unique spectra (non-phosphorylated and phosphorylated) were compared so these changes may be driven by changes in the local environment altering H/D exchange, rather than strictly changes in secondary structure. No changes in TTC7B and FAM126A were observed (significant change defined as greater than both 5% and 0.4 Da at any time point in any peptide with an unpaired two-tailed t-test p<0.01).

### In the presence of EFR3 Y2090 phosphorylation does not alter membrane recruitment, but does inhibit lipid kinase activity

Although ^WT^PI4KA^pY1154,pY2090^ displayed a clear decrease in lipid kinase activity in the presence of 100% PI vesicles, this experimental system is not a good biological mimic of cellular PI4KA activity. *In vivo*, EFR3 is required for robust recruitment of PI4KA to the plasma membrane^7,9^. We reasoned that the inhibition observed in the absence of EFR3 could be due to altered membrane binding of the PI4KA complex, which might have a negligible effect under EFR3 mediated recruitment. First, we tested the ability of ^WT^PI4KA^pY1154,pY2090^ to bind to the C-terminus of EFR3A. We used biolayer interferometry (BLI) to measure the association of immobilised EFR3A to either ^WT^PI4KA^pY1154,pY2090^ or ^WT^PI4KA **(Fig. S8A**). The association and dissociation rates of these complexes were indistinguishable from each other, with values consistent with previously measured rates **(Fig. S8B**).

Next, we sought to measure PI4KA recruitment and kinase activity on supported lipid bilayers (SLBs) with lipid compositions that more closely resembled that of the plasma membrane in the presence of membrane-tethered EFR3. For these experiments, we used the SpyCatcher-SpyTag system to covalently attach the C-terminus of EFR3A to SLBs, as in a previous reconstitution of PI4KA membrane recruitment^10^ **(Fig. 5A**). To visualize dynamic membrane recruitment, we labeled PI4KA complexes with AF647. We then split a single batch of fluorescently tagged PI4KA-TTC7B-FAM126A complex into an unphosphorylated sample and a Lck-treated sample (AF647-^WT^PI4KA and AF647-^WT^PI4KA^pY1154,pY2090^, respectively). Based on MS analysis, we achieved ∼90% phosphorylation of residue Y2090 through Lck treatment (**source data**). In agreement with our BLI data, EFR3A-mediated bulk SLB recruitment of AF647-^WT^PI4KA was not greatly perturbed by phosphorylation of Y2090 (**Fig. S8C**). Single molecule dwell time analysis (**Fig. S8D**) showed a slight perturbation with Y2090 phosphorylation but with no changes in cumulative membrane binding frequency measurements (**Fig. S8E**). The modest decrease in the single molecule dwell times of AF647-^WT^PI4KA^pY1154,pY2090^ could be attributed to the inability of the kinase to efficiently engage the lipid substrate once bound to membrane tethered EFR3A.

**Figure 5.**
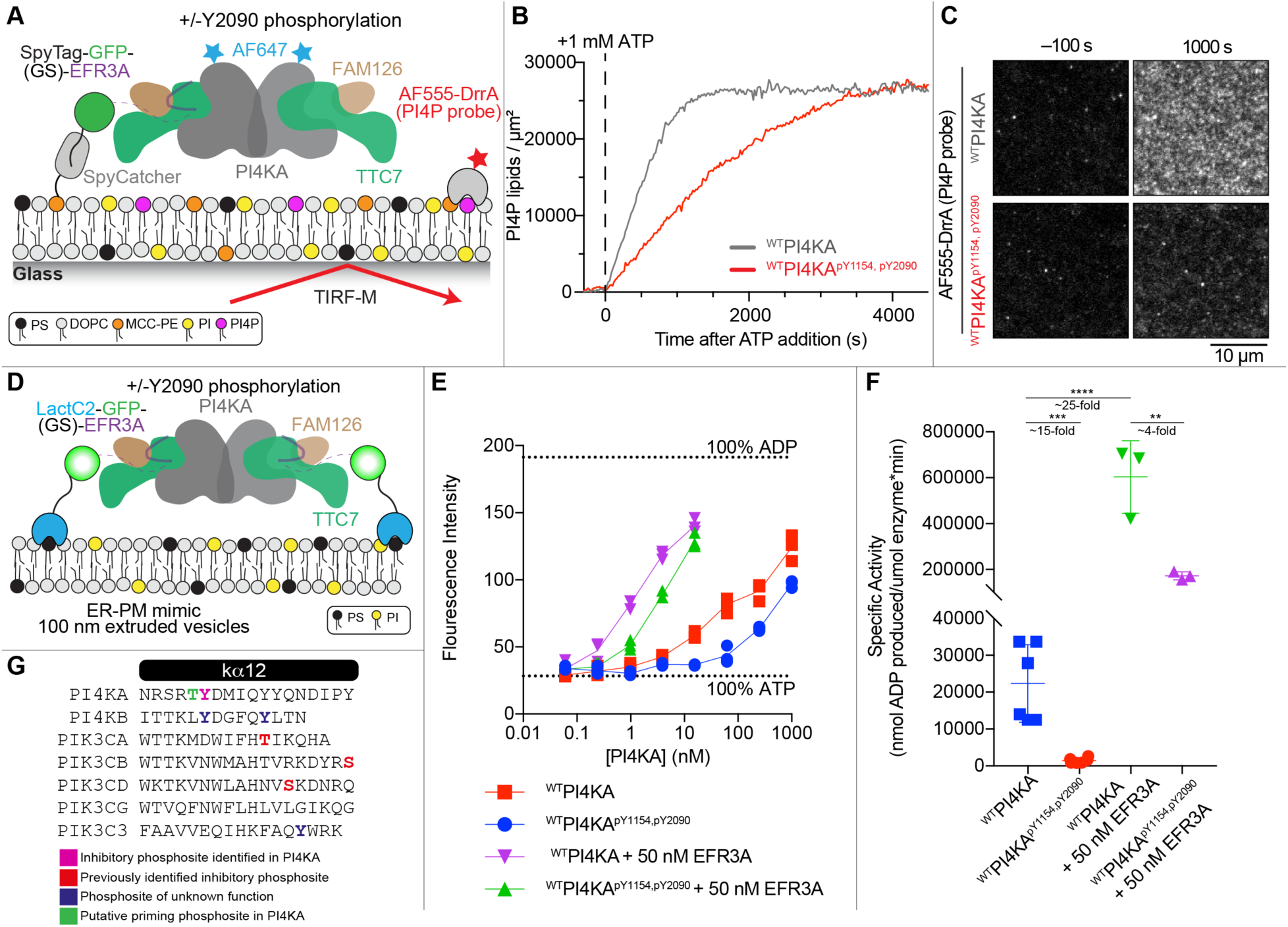
Y2090 phosphorylation inhibits PI4P production even when tethered to a membrane by EFR3. A. Cartoon schematic showing the experimental design for measuring AF647-PI4KA localisation and activity on an SLB containing tethered EGFP-SpyTag-EFR3A +/− phosphorylation of pY1154 and pY2090. SpyCatcher is covalently linked to a maleimide lipid (i.e. MCC–PE) and EGFP-SpyTag-EFR3A is conjugated to SpyCatcher via an isopeptide bond. A soluble AF555 labeled PI4P biosensor derived from DrrA is used to monitor PI4P production. Membrane localisation of EGFP-SpyTag-EFR3A, AF647– ^WT^PI4KA, and AF555-DrrA are visualized by TIRF microscopy. B. Phosphorylation attenuates PI4KA lipid kinase activity. The production of PI4P was monitored by quantifying membrane recruitment of 20 nM AF555-DrrA (PI4P probe) following the addition of 1 mM ATP. Lipid phosphorylation was measured using conditions in (Fig. S8F). C. Representative TIRF-M image showing the localisation of 20 nM AF555-DrrA in (H) before (−100 s) and after (1000 s) adding 1 mM ATP to stimulate AF647-^WT^PI4KA +/− phosphorylation lipid kinase activity. B-C. Membrane composition: 86% DOPC, 10% PS, 2% PI, 2% MCC-PE (SpyCatcher conjugated). D. Cartoon schematic showing the experimental design for in vitro membrane reconstitution of ^WT^PI4KA and ^WT^PI4KA^pY1154,pY2090^ onto EFR3A anchored 100 nm extruded ER-PM mimic vesicles (10% PI, 25% PS, 40% PE, 15% PC, 10% Cholesterol). EFR3A is anchored to the vesicles through an N-terminal LactC2 domain, which is an established phosphatidylserine biosensor_45_. E. Measurement of ATP turnover of ^WT^PI4KA and ^WT^PI4KA^pY1154,pY2090^ +/− 50 nM LactC2-GFP-GS-EFR3A on ER-PM mimic vesicles (10% PI, 25% PS, 40% PE, 15% PC, 10% Cholesterol). Experiments were performed with 0.06-1000 nM and a final vesicle concentration of 0.5 mg/ml. (Technical replicates; individual data points are shown, n=3) F. Specific activity values for ^WT^PI4KA and ^WT^PI4KA^pY1154,pY2090^ +/− 50 nM LactC2-GFP-GS-EFR3A on ER-PM mimic vesicles (10% PI, 25% PS, 40% PE, 15% PC, 10% Cholesterol). Specific activity was calculated using the concentrations where kinase activity was in range (3.91 and 15.63 nM for ^WT^PI4KA, 62.5 and 250 nM for ^WT^PI4KA^pY1154,pY2090^, 0.24 nM for ^WT^PI4KA + 50 nM EFR3A, and 0.98 nM for ^WT^PI4KA^pY1154,pY2090^ + 50 nM EFR3 (Technical replicates; data are presented as mean values, and error bars are SD, n=3 for each concentration). Two-tailed t-test P values are shown. G. Multiple sequence alignment comparing the kα12 helix across human PI4K and PI3K enzymes with previously identified phosphosites and putative phosphosites.

To determine whether phosphorylation of PI4KA modulates lipid kinase activity in the context of EFR3-mediated membrane recruitment, we used an AF555 labeled PI4P biosensor derived from DrrA to monitor the production of PI4P on SLBs. This allowed us to simultaneously quantify PI4KA membrane localisation and catalytic activity for both phosphorylated and non-phosphorylated PI4KA complexes. After allowing AF647-PI4KA to equilibrate on an SLB through interactions with membrane-tethered EFR3A, lipid kinase activity measurements were initiated by adding 1 mM ATP (**Fig. 5B**). Although similar membrane densities of ^WT^PI4KA and ^WT^PI4KA^pY1154,pY2090^ were achieved (**Fig. S8F**), ^WT^PI4KA^pY1154,pY2090^ exhibited a notable decrease in lipid kinase activity (**Fig. 5B)**, as indicated by decreased membrane density of DrrA (**Fig. 5C**).

In tandem, we established an *in vitro* kinase activity assay to analyze the effect of Y2090 phosphorylation downstream of EFR3 recruitment where we reconstituted EFR3A’s C-terminus onto Endoplasmic Reticulum (ER)-PM mimic 100 nm vesicles (10% PI, 25% PS, 40% PE, 15% PC, 10% Cholesterol). We anchored the C-terminus of EFR3 onto membranes through attachment of an N-terminal LactC2 domain, which is a high affinity phosphatidylserine binder^45^, in place of the SpyTag (**Fig. 5D)**. In agreement with kinase assays on 100% PI vesicles, we found that ATP-turnover in the presence of ER-PM vesicles was reduced by ∼15-fold in ^WT^PI4KA^pY1154,pY2090^, compared to ^WT^PI4KA (**Fig. 5E/F**). The presence of EFR3 dramatically increased PI4KA lipid kinase activity, with a > 25-fold increase (**Fig. 5E/F**). PI4KA activity was still inhibited by Y2090 phosphorylation downstream of EFR3 recruitment, with this being ∼4 fold less (**Fig. 5E/F**). These results demonstrate that even when PI4KA is bound to membrane-tethered EFR3, Y2090 phosphorylation attenuates PI4KA lipid kinase activity.

## Discussion

Essential to PI4KA’s function is its targeted recruitment to the plasma membrane, where it sustains the PI4P pool that mediates phosphatidylserine transport, and acts as the precursor molecule for PI(4,5)P_2_ and PI(3,4,5)P_3_ generation that are required for calcium and Akt mediated signalling cascades. PI4P depletion at the plasma membrane through inhibition of PI4KA does not have a major effect on the abundance of PI(4,5)P_2_^46–48^, however, PI4KA inhibition does dramatically alter oscillations of PI4P and PI(4,5)P_2_ which seem to play important roles in controlling signalling downstream of these molecules^14^. This seems to indicate a critical role of PI4P flux in mediating signalling events downstream of PI4KA, making it crucial for us to understand how PI4KA activity is regulated. The most well understood regulator of PI4KA PM localisation and activity is the plasma membrane localised protein EFR3A/B which is anchored to the plasma membrane by palmitoylation^7^. We previously determined that the C-terminal tail of EFR3 engages with TTC7 and FAM126 to bring PI4KA to the plasma membrane, with deletion of this region abolishing PM recruitment^9,10^. However, our understanding of how PI4KA activity and EFR3 recruitment is regulated is still unknown. Multiple phosphatases have been identified that alter PI4KA activity, with the protein phosphatase calcineurin increasing PI4KA activity following PI(4,5)P₂ depletion^12,13^. Phosphorylation at Y723 of EFR3B in the TTC7-FAM126 interface decreases EFR3B’s interaction with the PI4KA complex^10^, with the exact kinases and phosphatases regulating this being unidentified. This highlighted the potential for PI4KA activity to be controlled by phosphorylation, either directly or indirectly. Here we identified a direct inhibitory phosphorylation site in PI4KA’s kα12 helix at Y2090, that substantially decreases PI4KA activity.

PI4KA is evolutionarily conserved with PI4KB and the class I, II, and III PI3Ks, with a conserved catalytic architecture composed of a helical domain and a bi-lobal kinase domain^16^. These all share a similar regulatory motif in their kinase domain composed of helices kα7–kα12, that surround the activation loop and play a key role in membrane association. The final C-terminal helix kα12 is absolutely essential for membrane binding, as truncation of this region completely inactivates lipid kinase activity for PI4KB, PIK3CA, PIK3CB, and PIK3C3^17,49–51^. Recent cryo-EM analysis of the PIK3CA enzyme interacting with nanodiscs reveal the likely mechanistic basis for the importance of this C-terminal helix, with the amphipathic helix inserting its hydrophobic face into the membrane plane^20^.

Intriguingly, multiple studies have reported inhibitory phosphorylation sites in the kα12 helix of class I PI3Ks (**Fig. 5G**). PIK3CA can be phosphorylated downstream of the hippo kinases at T1061, with phosphorylation dramatically decreasing both lipid kinase activity and recruitment to negatively charged membranes^23^. Both PIK3CB and PIK3CD have been reported to auto-phosphorylate kα12 residues (S1070 in PIK3CB, and S1039 in PIK3CD)^21,22^, leading to decreased lipid kinase activity. Intriguingly, both Vps34 (PIK3C3) and PI4KB have phosphosites in kα12 that have been reported (Y884 in Vps34 and Y805 in PI4KB), but the effect of these PTMs have not been characterised. However, a Y884D-W885D mutant of Vps34 almost completely inhibits lipid kinase activity^52^, suggesting that Y884 phosphorylation would be inhibitory. The putative mechanism of inhibition for all these sites is that addition of a negatively charged residue in the kα12 helix causes charge-charge repulsion with negatively charged membranes, preventing kα12 membrane association. This mechanism is consistent with our data comparing PI4KA lipid kinase activity of phosphorylated PI4KA with and without membrane localised EFR3. These experiments showed that Y2090 phosphorylation caused a larger decrease in activity in the absence of EFR3 (∼20 fold), with a smaller decrease when membrane localised EFR3 was present (∼4 fold), as EFR3 can partially overcome decreased membrane association caused by kα12 phosphorylation. Overall, this supports kα12 phosphorylation as a conserved mode of inhibition across multiple PI4Ks and PI3Ks.

While our work identifies an inhibitory phospho-site in the C-terminus of PI4KA, an important outstanding question is what upstream signalling pathways engage Y2090 phosphorylation *in vivo*, and under what cellular conditions. Our initial focus on Y1154, the most frequently annotated site in phosphoproteomic databases with over 250 reported observations, ultimately proved instrumental in uncovering Y2090 regulation, despite Y1154 itself having no detectable effect on PI4KA kinase activity or stability. The weak conservation of Y1154 (restricted to higher mammals) and its maximum *in vitro* phosphorylation of ∼50%, suggesting modification of only one protomer in the PI4KA homodimer, suggest it may not represent a functionally important regulatory site. Further analysis of Y2090 in phosphoproteomic databases uncovered multiple additional identifications of Y2090 phosphorylation in multiple cell lines and primary tissues^33–36,53^. Overall, Y2090 is strictly conserved across all eukaryotes examined, is phosphorylated with high efficiency (∼90%) and causes significant decreases in lipid kinase activity, properties fully consistent with a bona fide regulatory phospho-switch.

A key mechanism by which tyrosine kinase signalling is amplified is through priming phosphorylation, wherein phosphorylation of a residue immediately upstream of a target tyrosine dramatically increases the efficiency of subsequent tyrosine phosphorylation^29^. Immediately N-terminal to Y2090 is a threonine at position T2089 (**Fig. 5G**), which has been found to be phosphorylated in phosphoproteomic databases^15^. When kinase predictions are run for pY2090 with priming phosphorylation at T2089, the score ranks of a broad spectrum of growth factor and immune receptor tyrosine kinases increase dramatically. Multiple members of the INSR, EPHR, FGFR, SRC, and VEGFR families enter the top 25, with the insulin receptor-related kinase (IRR) ranking first (**Table S2**). This is biologically compelling: many of these kinases are well-established activators of PI3K/Akt signalling, a pathway that is itself dependent on the PI4P-derived lipid PIP_2_ as its substrate precursor. This raises the hypothesis that Y2090 phosphorylation could constitute a previously unrecognised negative feedback loop in which activation of growth factor receptor tyrosine kinases inhibits PI4KA activity and limits the availability of the PIP_2_ substrate pool for continued PI3K signalling. Such a mechanism would represent an elegant means of self-limiting pro-growth signalling at the level of membrane lipid supply, and could additionally alter the broader plasma membrane environment through decreased PS density and/or Ras nanoclustering^2,27,28^. The priming phosphorylation model also offers a potential explanation for why pY2090 is rarely detected in standard phosphoproteomic screens, as co-phosphorylation of T2089 and Y2090 on the same peptide would complicate database matching and fragment ion assignment in typical data-dependent acquisition workflows.

A key limitation of our study is that the physiological context driving Y2090 phosphorylation remains undefined. Given PI4KA’s membrane localization, the local cellular environment likely dictates which tyrosine kinase mediates this modification. Our data showing that Y2090F mutations retain lipid kinase activity will allow for future cellular and *in vivo* analysis of the potential role of Y2090 phosphorylation feedback in a variety of immune and pro-growth conditions. Importantly, the identification of this PI4KA regulatory mechanism will be important in future pharmacological modulation of PI4KA, offering new opportunities to control its activity in both normal and disease-relevant settings.

## Experimental Procedures

### Multiple Sequence Alignments

Multiple sequence alignments were generated using Clustal Omega. Aligned sequences were then analyzed by ESPript 3.0 (https://espript.ibcp.fr)^54^. The uniprot accession codes for Figure 1 are P42356, E9Q3L2, A0A8M3AWF4, S4RUP9, V4C7L9, Q9W4X4, Q9XW63, Q9USR3, and P37297. The uniprot accession codes for Figure 5 are P42356, O00329, P42336, P42338, Q8NEB9, Q9UBF8, and P48736.

### Cloning

Plasmids containing genes for full-length SSH-PI4KA-TTC7B-FAM126A (1-308), referred to as ^WT^PI4KA were cloned into the MultiBac vector, pACEBac1 (Geneva Biotech) as described in^6^. Plasmids containing genes for full-length SSH-ybbr-PI4KA-TTC7B-FAM126A (1-308), referred to as ybbr-^WT^PI4KA, SSH-PI4KA Y1154F-TTC7B-FAM126A (1-308), referred to as ^Y1154F^PI4KA and SSH-PI4KA Y2090F-TTC7B-FAM126A (1-308), referred to as ^Y2090F^PI4KA were cloned in the pBIG1a biGBac vector using the biGbac cloning method ^55^. The kinase domain of Lck (225-509) was synthesized (ThermoFisher Gene Art Gene Synthesis) in a PET151 backbone with an N-terminal cleavable twin-strep, 6x his tag. For expression in *Sf9* cells residues the Lck (225-509) kinase domain was cloned into a pLIB vector with a cleavable Twin-strep tag and 10x his. INSR (1011-1382) and SRC FL were synthesized (ThermoFisher Gene Art Gene Synthesis) into a FastBac backbone with N-terminal cleavable twin-strep and 10x his tags. EFR3A (721 to 791) was subcloned into a pOPT vector containing an N-terminal MBP tag in addition to a cleavable Twin-strep tag and 10x his. GFP-SpyTag-(GS)-EFR3A (721-791) was cloned into a pOPT vector with N-terminal 2x strep tag, 10x his tag and TEV cleavage site followed by the EGFP-Spytag with a GS linker before residues 721-791 of EFR3A (Referred to as EGFP-SpyTag-(GS)-EFR3A). LactC2-EGFP-EFR3A (721-791) was synthesized (ThermoFisher Gene Art Gene Synthesis) into a pOPT vector containing a cleavable Twin-strep tag and 10x his. Final pACEbac1, pBIG1a, FastBac and pLIB constructs were transformed into DH10EmBacY cells (Geneva Biotech), with white colonies indicating successful generation of bacmids containing our genes of interest. Full list of all plasmids used are shown in **Table S3**.

### Protein Expression

Bacmid harbouring ^WT^PI4KA, ybbr-^WT^PI4KA, ^Y1154F^PI4KA, ^Y2090F^PI4KA, Lck (225-509), SRC (FL), and INSR (1011-1382) were transfected into *Spodoptera frugiperda (Sf9) cells*, and viral stocks were amplified for one generation to acquire a P2 generation final viral stock. Final viral stocks were added to *Sf9* cells in a 1/100 virus volume to cell volume ratio. Constructs were expressed for 65-72 hours before harvesting the infected cells.

Plasmid containing the coding sequences for Lck (225-509), EGFP-SpyTag-EFR3A, and LactC2-EGFP-EFR3A were expressed in BL21 DE3 C41 *Escherichia coli* and induced with 0.1 mM IPTG and grown at 18°C overnight. MBP-EFR3A (721-791) was induced with 0.5 mM IPTG and grown at 37°C for 3 hours. All cell pellets were washed with PBS prior to flash freezing and storage at −80°C.

### Protein Purification

#### ^WT^PI4KA, ybbr-^WT^PI4KA, ^Y1154F^PI4KA, and ^Y2090F^PI4KA protein purification

*Sf9* pellets were resuspended in lysis buffer [20 mM imidazole pH 8.0, 100 mM NaCl, 5% glycerol, 2 mM βMe, protease (Protease Inhibitor Cocktail Set III, Sigma)] and lysed by sonication. Triton X-100 was added to 0.1% v/v final, and lysate was centrifuged for 45 min at 20,000×g at 1 °C. The supernatant was then loaded onto a 5 ml HisTrap column (Cytiva) that had been equilibrated in nickel–nitrilotriacetic acid (Ni–NTA) A buffer (20 mM Tris [pH 8.0], 100 mM NaCl, 20 mM imidazole [pH 8.0], 5% [v/v] glycerol, and 2 mM bME). The column was washed with 3-4 CV of Ni-NTA A buffer (20 mM imidazole [pH 8.0], 100 mM NaCl, 5% [v/v] glycerol, and 2 mM bME) followed by 3-4 CV of 6% Ni–NTA B buffer (27 mM Imidazole [pH 8.0], 100 mM NaCl, 5% [v/v] glycerol, and 2 mM bME) before being eluted with 3-4 CV of 100% Ni–NTA B. Eluted protein was loaded onto a 5 ml StrepTrapHP column (Cytiva) pre-equilibrated GFB (20 mM imidazole pH 7.0, 150 mM NaCl, 5% glycerol [v/v], 0.5 mM TCEP). The His-strep tag was cleaved with Lip-TEV with cleavage proceeded overnight. Cleaved protein was eluted with GFB and concentrated in a 50 kDa MWCO concentrator (MilliporeSigma). See Fluorescent labeling of ybbr-PI4KA-TTC7B-FAM126A for remainder of ybbr-^WT^PI4KA purification. For experiments directly comparing phosphorylated PI4KA to non-phosphorylated protein (HDX-MS and kinetic assays), the concentrated protein was split into two groups. PI4KA was treated with 0.1-0.18 mg/ml final of Lck (225-509), 1 mM final ATP, and 5 mM MgCl_2_ and left to incubate on ice for 3 hours to generate phosphorylated PI4KA. Non-phosphorylated PI4KA was treated equivalently, replacing Lck (225-509) with Lck buffer (20 mM HEPES pH 7.5, 150 mM NaCl, 0.5 mM TCEP). Concentrated protein was loaded onto the Superose 6 Increase 10/300 GL (Cytiva) pre-equilibrated in GFB. Protein fractions from a single peak were collected and concentrated in 50 kDa MWCO concentrator (Millipore Sigma), flash-frozen in liquid nitrogen and stored at −80 °C until further use. Phosphorylation identification and quantification was done using Mass Spectrometry, described below. ^WT^PI4KA^pY1154,pY2090^ was ∼50% phosphorylated at Y1154 and ∼90% phosphorylated at Y2090. ^Y1154F^PI4KA^pY2090^ was ∼95% phosphorylated at Y2090 and ^Y2090F^PI4KA^pY1154^ was ∼40% phosphorylated at Y1154, with Y2096/2102 being ∼20% phosphorylated only in the Y2090F mutant.

#### Lck (225-509), INSR (1011-1382), and SRC purification

Cell pellets were lysed by sonication for 5 min in lysis buffer (20 mM Tris [pH 8.0], 100 mM NaCl, 5% [v/v] glycerol, 20 mM imidazole, 2 mM β-mercaptoethanol [bME]), and protease inhibitors [Millipore Protease Inhibitor Cocktail Set III, EDTA free]). Triton X-100 was added to 0.1% v/v, and the solution was centrifuged for 45 min at 20,000*g* at 1 °C (Beckman Coulter J2-21, JA-20 rotor). The supernatant was then loaded onto a 5 ml HisTrap column (Cytiva) that had been equilibrated in nickel–nitrilotriacetic acid (Ni–NTA) A buffer (20 mM Tris [pH 8.0], 100 mM NaCl, 20 mM imidazole [pH 8.0], 5% [v/v] glycerol, and 2 mM βME). The column was washed with 4 CV of Ni–NTA A buffer, followed by 4 CV of 6% Ni–NTA B buffer (20 mM Tris [pH 8.0], 100 mM NaCl, 200 mM imidazole [pH 8.0], 5% [v/v] glycerol, and 2 mM βME) before being eluted with 4 CV of 100% Ni–NTA B. The eluate was then loaded onto a 5 ml StrepTrapHP column (Cytiva) and washed with 2.5 CV of GFB (20 mM HEPES [pH 7.5], 150 mM NaCl, and 0.5 mM Tris(2-carboxyethyl) phosphine [TCEP]. The His-strep tag was cleaved from Lck (225-509) and INSR (1011-1382) with strep tagged Lip-TEV with cleavage proceeded overnight. Cleaved protein was eluted with 2 CV of GFB. Src was eluted with 4 CV of GFB containing 2.5 mM desthiobiotin and concentrated in a 30 kDa MWCO concentrator (Millipore Sigma). Concentrated proteins were treated with 1 mM ATP and 2 mM MgCl_2_ for 1 minute at RT and then loaded onto the Superdex 75 Increase 10/300 GL (Cytiva) (Lck (225-509) and INSR (1011-1382)) or the Superdex 200 Increase 10/300 GL (Cytiva) (SRC) pre-equilibrated in GFB. Protein fractions were collected and concentrated in a 10 kDa MWCO concentrator (Millipore Sigma), flash frozen in liquid nitrogen, and stored at −80 °C.

#### MBP-EFR3A, EGFP-SpyTag-EFR3A, and LactC2-EGFP-EFR3A purification

Cell pellets were lysed by sonication for 5 min in lysis buffer (20 mM Tris [pH 8.0], 100 mM NaCl, 5% [v/v] glycerol, 20 mM imidazole, 2 mM β-mercaptoethanol [bME]), and protease inhibitors [Millipore Protease Inhibitor Cocktail Set III, EDTA free]). Triton X-100 was added to 0.1% v/v, and the solution was centrifuged for 45 min at 20,000*g* at 1 °C (Beckman Coulter J2-21, JA-20 rotor). The supernatant was then loaded onto a 5 ml HisTrap column (Cytiva) that had been equilibrated in nickel–nitrilotriacetic acid (Ni–NTA) A buffer (20 mM Tris [pH 8.0], 100 mM NaCl, 20 mM imidazole [pH 8.0], 5% [v/v] glycerol, and 2 mM βME). The column was washed with 4 CV of high salt buffer (20 mM Tris [pH 8.0], 1 M NaCl, 5% [v/v] glycerol, and 2 mM βME), then 3-4 CV of Ni–NTA A buffer, followed by 3-4 CV of 6% Ni–NTA B buffer (20 mM Tris [pH 8.0], 100 mM NaCl, 200 mM imidazole [pH 8.0], 5% [v/v] glycerol, and 2 mM βME) before being eluted with 3-4 CV of 100% Ni–NTA B. The eluate was then loaded onto a 5 ml StrepTrapHP column (Cytiva) and washed with 3-4 CV of high salt gel filtration buffer (GFB) (20 mM HEPES [pH 7.5], 500 mM NaCl, 5% [v/v] glycerol and 1 mM Tris(2-carboxyethyl) phosphine [TCEP]), then 1 CV of GFB containing 2 mM ATP, 10 mM MgCl_2_, and 150 mM KCl, followed by 4 CV of GFB (20 mM HEPES [pH 7.5], 150 mM NaCl, 5% [v/v] glycerol and 1 mM Tris (2-carboxyethyl) phosphine [TCEP]). MBP-EFR3A protein was eluted with 3 CV of GFB containing 2.5 mM desthiobiotin and concentrated in a 10 kDa MWCO concentrator (Millipore Sigma). The his-strep tag of the EGFP-SpyTag-EFR3A was removed overnight at 4°C using tobacco etch virus protease containing a stabilizing lipoyl domain (Lip-TEV). Concentrated protein was loaded onto the Superdex 200 Increase 10/300 GL (Cytiva) pre-equilibrated in GFB. Protein fractions were collected and concentrated in a 10 kDa MWCO concentrator (Millipore Sigma), flash frozen in liquid nitrogen, and stored at −80 °C. LactC2-EGFP-EFR3A was purified in the same way (uncleaved); however, was loaded onto a StrepTrap XT column and eluted with 4 CV of elution buffer containing 50 mM biotin, followed by elution with an additional 3 CV mL with 50 mM Biotin and 0.1 mM EDTA. Additionally, all buffers included 300 mM NaCl, with the NiNTA buffers containing 10% Glycerol.

#### KCK-SpyCatcher

A his_10_-TEV-SUMO-KCK-SpyCatcher fusion was expressed and purified from Rosetta(DE3) *E. coli*. Bacteria were grown at 37°C in Terrific Broth until reaching OD_600_ = 0.8. Cultures were shifted to 18°C for 1 hour then induced with 0.1 mM IPTG. Expression was allowed to continue for 20 hours before harvesting. Cells were lysed in 50 mM Na_2_HPO_4_ [pH 8.0], 400 mM NaCl, 10 mM imidazole, 4 mM BME, 1 mM PMSF, 100 μg/mL DNase by sonication. Lysate was centrifuged at 15,000 rpm (35,000 x *g*) for 60 minutes in a Beckman JA-20 rotor at 4°C. NiNTA resin was added to the supernatant and allowed to bind for 2 hours, then washed with 50 mM Na_2_HPO_4_ [pH 8.0], 400 mM NaCl, 30 mM imidazole, 4 mM BME. Bound protein was eluted in 50 mM Na_2_HPO_4_ [pH 8.0], 400 mM NaCl, 500 mM imidazole, 4 mM BME. Peak fractions were pooled and loaded onto a desalting column equilibrated in 50 mM Na_2_HPO_4_ [pH 8.0], 400 mM NaCl, 20 mM imidazole, 5% glycerol, 1 mM BME to remove most of the imidazole. A his_6_-SenP2 protease was added to the eluted protein which was then stored overnight at 4°C. The cleaved protein was recirculated over NiNTA resin to capture the tag and protease. The flowthrough was concentrated in a 10 kDa MWCO Amicon-Ultra concentrator before loading onto a 124 mL Superdex 75 column equilibrated in 20mM Tris [pH 8.0], 200 mM NaCl, 10% glycerol, 1 mM TCEP. Peak fractions were pooled, concentrated to a minimum concentration of 150 µM, and snap frozen in liquid nitrogen for storage at −80°C.

#### DrrA

The PI(4)P lipid binding domain, DrrA(544-647aa), was expressed in BL21(DE3) *E. coli* as a his_6_-MBP-TEV-GGGGG-DrrA(544-647aa) fusion as previously described ^56^. Bacteria were grown at 37°C in Terrific Broth until it reached OD_600_ = 0.8. Cultures were shifted to 18°C for 1 hour then induced with 0.1 mM IPTG. Expression was allowed to continue for 20 hours before harvesting. Cells were lysed in 50 mM Na_2_HPO_4_ [pH 8.0], 300 mM NaCl, 0.4 mM BME, 1 mM PMSF, 100 μg/mL Dnase using a microfluidizer. Lysate was centrifuged at 16,000 rpm (35,172 x *g*) for 60 minutes in a Beckman JA-17 rotor at 4°C. Lysate was circulated over a 5 mL HiTrap Chelating column (GE Healthcare, Cat# 17-0409-01) charged with CoCl_2_. Bound protein was eluted with a linear gradient of imidazole (0-500 mM). TEV protease was added peak fractions to cleave off the his_6_-MBP tag. The TEV cleavage was performed while dialyzing the partially pure DrrA for 16 hours at 4°C against 4 liters of buffer containing 50 mM Na_2_HPO_4_ [pH 8.0], 300 mM NaCl, and 0.4 mM BME. The next day, dialysate containing cleaved protein was recirculated over a 5 mL HiTrap Chelating column to capture the his_6_-MBP and his_6_-TEV. Flowthrough containing GGGGG-DrrA(544-647aa) was concentrated using a 5 kDa MWCO Vivaspin 20 before being loaded on a Superdex 75 (10/300 GL) size exclusion column equilibrated in 20 mM Tris [pH 8.0], 200 mM NaCl, 10% glycerol, 1 mM TCEP. Peak fractions were pooled and snap frozen in liquid nitrogen for storage at −80°C.

Sortase mediated peptide ligation was used to label GGGGG-DrrA(544-647aa) with AlexaFluor555 (AF555) as previously described ^56^. In brief, the sortase labeling reaction was prepared by combining 75 μM AF555-LPETGG, 28 μM GGGGG-DrrA(544-647aa), and 5 μM NusA-SortaseA in buffer containing 50 mM Tris [pH 8.0], 150 mM NaCl, 6 mM CaCl_2_, 1 mM TCEP. The labeling reaction was incubated at 18°C for 16 hrs. Unreacted AF555-LPETGG was mostly removed using a Nap25 G25 Sephadex desalting column. The remaining AF555-LPETGG and NusA-SortaseA were separated from AF555-LPETGGGGG-DrrA(544-647aa) using a Superdex 75 size exclusion column equilibrated in 20 mM Tris [pH 8.0], 200 mM NaCl, 10% glycerol, 1 mM TCEP. Peak fractions were pooled and snap frozen in liquid nitrogen for storage at −80°C.

#### NusA-SortaseA

A NusA-his_6_-SortaseA fusion was expressed in BL21(DE3) *E. coli*. Bacteria were grown at 37°C in Terrific Broth until reaching OD_600_ = 0.8. Cultures were shifted to 18°C for 1 hour then induced with 0.1 mM IPTG. Expression was allowed to continue for 20 hours before harvesting. Cells were lysed in 50 mM Na_2_HPO_4_ [pH 8.0], 400 mM NaCl, 4 mM BME, 1 mM PMSF, 100 μg/mL Dnase by sonication. Lysate was centrifuged at 15,000 rpm (35,000 x *g*) for 60 minutes in a Beckman JA-20 rotor at 4°C.

NiNTA resin was added to the supernatant and allowed to bind for 2 hours, then washed with 50 mM Na_2_HPO_4_ [pH 8.0], 400 mM NaCl, 10 mM imidazole, 4 mM BME. Bound protein was eluted in 50 mM Na_2_HPO_4_ [pH 8.0], 400 mM NaCl, 500 mM imidazole, 4 mM BME. Peak fractions were pooled and dialyzed against 4 liters of buffer containing 20 mM Tris [pH 8.0], 200 mM NaCl, 5 mM BME. Dialysate was concentrated in a 10 kDa MWCO Amicon-Ultra concentrator before loading onto a 124 mL Superdex 75 column equilibrated in 20 mM Tris [pH 8.0], 150 mM NaCl, 10% glycerol, 1 mM TCEP. Peak fractions were pooled and snap frozen in liquid nitrogen for storage at −80°C. Note that the NusA solubility tag was intentionally left attached to sortase to increase the molecular weight to 75 kDa, which made it easier to separate from the DrrA(544-647aa) after labeling with AF555.

### Preparation of supported lipid bilayers

Supported lipid bilayers (SLBs) were created to maximize membrane fluidity at room temperature (i.e. 23°C). The following lipids were used to generate small unilamellar vesicles (SUVs) by hydration and extrusion: 1,2-dioleoyl-sn-glycero-3-phosphocholine (18:1 DOPC, Avanti #850375C), 1,2-dioleoyl-sn-glycero-3-phospho-L-serine (18:1 DOPS, Avanti #840035C), 1,2-dioleoyl-sn-glycero-3-phosphoethanolamine-N-[4-(p-maleimidomethyl)cyclohexane-carboxamide] (18:1 MCC-PE, Avanti #780201C), and phosphatidylinositol from bovine liver (PI, Avanti #840042C). To prepare the small unilamellar vesicles, a total of 2 µmoles lipids were combined in a 35 mL glass round bottom flask containing ∼2 mL of chloroform. Lipids were dried to a thin film using rotary evaporation with the glass round-bottom flask submerged in a 42°C water bath. After evaporating all the chloroform, the round bottom flask was place under a vacuum overnight at 23°C. The lipid film was then resuspended in 2 mL of 0.45 µm filtered 1x PBS pH 7.4 (Corning, cat# 46-013-CM), bring the lipid suspension to a final concentration of 1 mM. To generate 50 nm SUVs, the 1 mM total lipid mixtures was extruded through a 0.05 µm pore size 19 mm polycarbonate (PC) membrane (Cytiva Whatman, Cat# 800308) with filter supports (Cytiva Whatman, Cat# 230300) on both sides of the PC membrane. All lipid compositions used for supported lipid bilayer experiments contained the following molar percentages: 86% DOPC, 10% PS, 2% PI, 2% MCC-PE.

Supported lipid bilayers were formed on 25 x 75 mm coverglass (IBIDI, #10812) as previously described ^57^. In brief, coverglass was first cleaned with 2% Hellmanex III (ThermoFisher, Cat#14-385-864) and then etched with Piranha solution (1:3, hydrogen peroxide:sulfuric acid) for 15 minutes. Etched coverglass, washed extensively with MilliQ water again, is rapidly dried with nitrogen gas before adhering to a 6-well sticky-side chamber (IBIDI, Cat# 80608). SLBs were formed by flowing 0.25 mM total lipid concentration of 50 nm SUVs diluted in 1x PBS pH [7.4] into an assembled IBIDI chamber. SUVs were incubated in the IBIDI chamber for 30 minutes and then washed with 4 mL of PBS [pH 7.4] to remove non-absorbed SUVs. Membrane defects are blocked for 5 minutes with a solution of clarified 1 mg/mL beta casein (ThermoFisher, Cat# 37528) in 1x PBS [pH 7.4]. After blocking membranes with 1 mg/mL beta casein for 5 minutes, bilayers were washed with 4 mL of 1x PBS. Reaction/imaging buffer was exchanged into the chamber prior to each experiment.

Conjugation of SpyCatcher to supported membranes was achieved as previously described^58^. Supported membranes containing 2% MCC-PE lipids were used to covalently couple SpyCatcher containing single N-terminal cysteine (i.e. thiol). For membrane coupling, 100 µL of 20 µM SpyCatcher was diluted in a 1x PBS [pH 7.4] and 0.1 mM TCEP buffer. This was incubated in an IBIDI chamber containing the SLB for 2 hours at 23°C. Unreacted MCC-PE lipids were then quench for 15 minutes 5 mM beta-mercaptoethanol (BME) diluted in 1x PBS [pH 7.4]. Quenched membranes were then washed and stored in 1x PBS [pH 7.4] until ready for imaging. To attach EGFP-SpyTag-EFR3A, a solution containing 100 nM protein was flowed into the chamber containing SpyCatcher membrane conjugated to MCC-PE lipids. After 20 minutes of SpyTag conjugation to SpyCatcher, unbound SpyTag proteins were flushed from the chamber with 1x PBS [pH 7.4]. These membranes were subsequently used to visualize EFR3A-mediated membrane recruitment of AF647-PI4KA.

### Fluorescent labeling of ybbr-PI4KA-TTC7B-FAM126A

The AF647-coenzyme A (CoA) derivative was generated in-house by combining 15 mM AF647 C_2_ Maleimide (Invitrogen, no. A20347) in dimethyl sulfoxide with 10 mM CoA (Sigma-Aldrich, no. C3144, MW = 785.33 g/mol) dissolved in 1× PBS to a final concentration of 7.5 mM AF647 C_2_ maleimide and 5 mM CoA. This mixture was incubated overnight at 20°C in an amber tube. AF647-CoA conjugate was wrapped in parafilm and stored at −20°C. We labeled recombinant ^WT^PI4KA-TTC7B-FAM126A (1-308) containing an N-terminal ybbR13 motif (DSLEFIASKLA) using Sfp transferase and AF647-CoA ^59^. Prior to the labelling of the protein, 10 μM AF647-CoA and 5 mM DTT in GFB (20 mM imidazole pH 7.0, 150 mM NaCl, 5% glycerol, 0.5 mM TCEP) were combined and incubated for 5 minutes at 20°C followed by 2 minutes on ice. Subsequently, chemical labeling was achieved by adding ybbr-^WT^PI4KA-TTC7B-FAM126A (1-308) (final 8 μM), Sfp-his6 (final 5 μM), and 10 mM MgCl_2_ and incubating the reaction overnight on ice. Excess AF647-CoA was removed using a HiTrap® desalting column preequilibrated in 3CV of GFB. Protein was then split into two groups and treated similarly to above (see protein purification) to generate AF647-^WT^PI4KA and AF647-^WT^PI4KA^pY1154,pY2020^. The proteins were then concentrated in a 50 kDa MWCO concentrator (Millipore Sigma) and loaded on a Superose 6 Increase 10/300 GL (Cytiva) equilibrated in GFB. Protein fractions were collected and concentrated in 50 kDa MWCO concentrator (Millipore Sigma), flash-frozen in liquid nitrogen and stored at −80 °C until further use. MS/MS samples were made containing 30 pmol of protein for phosphorylation quantification. Phosphorylation was quantified by generating extracted ion chromatograms of unphosphorylated and phosphorylated peptides with the percentage of phosphorylation being determined by dividing the area under the phosphorylated EIC by the sum of the phosphorylated and unphosphorylated EICs. Due to phosphorylation of Y1154 showing no decrease in activity in our lipid kinase assays, we only quantified phosphorylation of the Y2090 site in AF647-^WT^PI4KA^pY1154,pY2090^, which was ∼90%.

### TIRF Microscopy

All supported lipid bilayer TIRF-M experiments were performed using the following reaction buffer: 20 mM HEPES [pH 7.0], 150 mM NaCl, 1 mM ATP (Sigma, Cat #A2383-10G), 5 mM MgCl_2_, 0.5 mM EGTA, 20 mM glucose, 200 µg/mL beta casein (ThermoScientific, Cat #37528), 20 mM BME, 320 µg/mL glucose oxidase (Serva, Cat #22780.01 *Aspergillus niger*), 50 µg/mL catalase (Sigma, Cat #C40-100MG Bovine Liver), and 2 mM Trolox (Cayman Chemicals, Cat #10011659). For ATP spike-in experiments, ATP was initially omitted from the buffer and 1 mM ATP was spiked in at indicated timepoint. Perishable reagents (i.e. glucose oxidase, catalase, and Trolox) were added 5-10 minutes before acquiring images by TIRF-M. All reactions in this study were reconstituted and visualized on a TIRF microscope at 23°C.

Imaging experiments were performed using an inverted Nikon Ti2 microscope using a 100x (1.49 NA) oil immersion Nikon TIRF objective. The x-axis and y-axis positions were controlled using a Nikon motorized stage, joystick, and NIS element software. Fluorescently labeled proteins were excited with either a 561 or 637 nm OBIS laser diode (Coherent Inc. Santa Clara, CA) controlled with a Vortran laser launch (Sacramento, CA) and acousto-optic tuneable filters (AOTF) control. Excitation and emission light was transmitted through a multi-bandpass quad filter cube (C-TIRF ULTRA HI S/N QUAD 405/488/561/638; Semrock) containing a dichroic mirror. Fluorescence emission was detected on an iXion Life 897 EMCCD camera (Andor Technology Ltd., UK) after passing through a Nikon Ti2 emission filter wheel containing the following 25 mm emission filters: ET525/50M, ET600/50M, ET700/75M (Semrock). All experiments were performed at room temperature (23°C). Microscope hardware was controlled with Nikon NIS elements.

#### Single particle tracking

Single fluorescent PI4KA complexes bound to supported lipid bilayers were identified and tracked using the ImageJ/Fiji TrackMate plugin ^60^. As previously described ^57^, image stacks were loaded into ImageJ/Fiji as .nd2 files. Using the LoG detector, fluorescent particles were identified based on their size, brightness, and signal-to-noise ratio. The LAP tracker was used to generate trajectories that followed particle displacement as a function of time. FParticle trajectories were then filtered based on Track Start (rparticles at start of movie were removed), Track End (particles at end of movie were removed), Duration (particles present for less than 2 frames were removed), and Track displacement (immobilized particles were removed). The output files from TrackMate were then analyzed using Prism 9 graphing software.

To calculate membrane dwell times, cumulative distribution frequency (CDF) plots with the bin size set to the image acquisition frame interval (e.g. 52 ms) were created. The log_10_(1-cumulative distribution frequency) was plotted as a function of dwell time and fit to a single or double exponential curve as previously described ^57^. For the double exponential curve fits, the alpha value is the percentage of the fast-dissociating molecules characterized by the time constant, τ_1_. Data set contained dwell times measured for n≥1000 trajectories repeated as n=3 technical replicates.

#### Lipid phosphorylation kinetics

The production of PI(4)P lipids was monitored by TIRF-M using previously described methods ^56^. Reactions were reconstituted in the presence of 20 nM AF555-DrrA (PI(4)P biosensor) and 200 pM AF647-PI4KA (unphosphorylated and phosphorylated). ATP was initially omitted and the AF647-PI4KA was allowed to equilibrate with membrane tethered EFR3A for 15 minutes. Upon addition of 1 mM ATP, the density of membrane-bound AF555-DrrA increased linearly and then gradually plateaued. Under the assumption that no PI(4)P was present in the absence of ATP and that 2% PI lipids was converted to 2% PI(4)P at the plateau, the fluorescence value at the plateau was taken to represent 27,777 PI(4)P lipids/µm^2^.

### PI4KA treatment with tyrosine kinases

For the dose-response phosphorylation of ^WT^PI4KA by Lck (225-509), 2 uM was mixed with ATP (1 mM), GFB (20 mM HEPES [pH 7.5], 150 mM NaCl, 0.5 mM TCEP), MgCl_2_ (5 mM), and various amounts of Lck (225-509) (0 nM, 134 nM, 402 nM, 1200 nM, and 3600 nM). Reactions were incubated for 2 hours on ice and quenched with 60 µL of ice-cold acidic quench buffer (0.7 M guanidine-HCl, 1% formic acid), followed by immediate freezing using liquid nitrogen and storage at −80°C. This was done in independent triplicate (three unique reactions for each condition, with the same protein preparation of ^WT^PI4KA and Lck (225-509)). Additional samples of ^WT^PI4KA treated with either 0 ng or 1500 ng Lck were prepared in parallel, which, following freeze-thaw, were subjected to trypsin digestion and analyzed by LC–MS/MS on an Orbitrap Fusion using higher-energy collisional dissociation (HCD). The resulting MS/MS spectra were used to improve phosphosite localization and confirm the site of phosphorylation, as described in the Methods below.

For the experiment studying the ability of different kinases to phosphorylated Y1154 and Y2090, ^WT^PI4KA (2000 nM) was mixed with ATP (1 mM), GFB 150 mM NaCl, 5% Glycerol, 0.5 mM TCEP), MgCl_2_ (5 mM), and 1000 ng of Lck (225-509), INSR (1011-1382) or SRC for a final concentration of 3.02 μM, 1.5 μM, and 2.37 μM, respectively. An apo condition with no additional kinase was also generated. Reactions were incubated for 2 hr at on ice and quenched with 60 µL of ice-cold acidic quench buffer (0.7 M guanidine-HCl, 1% formic acid) followed by immediate freezing using liquid nitrogen and storage at −80°C.

Phosphorylation of all proteins was confirmed using mass spectrometry and FragPipe analysis. The LC-MS analysis of these samples was carried out using the same pipeline as used in the HDX-MS section. The loss of non-phosphorylated peptide ratios was determined by generating extracted ion chromatograms for each non-phosphorylated peptide using their molecular formula and charge state in the Bruker Compass Data Analysis software. The area under each extracted curve was then extracted. The only quantification that diverges from this method is the quantification of pY1154 and pY2090 in the cryo-EM specimen, and pY2090 in the AF647-^WT^PI4KA^pY1154,pY2090^, where the intensity of phosphorylated peptide was divided by the sum of phosphorylated and non-phosphorylated peptide intensities. This was done as there was not an untreated control sample to directly compare the non-phosphorylated peptide intensity. The full MS quantification of phosphorylated peptide abundance is provided in the **Source Data**.

### LC–MS/MS Analysis of Trypsin-Digested Samples

#### In-Solution Trypsin Digestion

To 7µg protein, 9 M urea/300 mM Tris pH 8.0/ 15 mM TCEP was added to samples for final concentration of 6 M urea, 10 mM TCEP and samples were incubated at 37°C for 30 minutes. Iodoacetamide (IAA) in MS-grade water was added to final concentration of 20 mM IAA and samples were incubated in the dark at room temperature for 30 minutes. The samples were diluted with 100 mM Tris pH 8.0 to reduce the urea concentration to 0.77 M. Modified sequencing grade porcine trypsin (Promega, V5111) was reconstituted in 100 mM Tris pH 8.0, trypsin was added to 50:1 substrate: enzyme ratio, and the samples were incubated at 37°C for 18 hours. The samples were acidified with formic acid (FA) to a final concentration of 1% FA, and desalted/concentrated using Pierce 100 µl C18 Tips (Thermo Scientific, catalogue number 87784) following the manufacturers protocol. The peptides were eluted in 50 µl 70% acetonitrile/0.1% FA and dried via speed vacuum. Samples were stored at −20°C until LC-MS/MS analysis.

#### LC-MS/MS analysis: Orbitrap Fusion

Samples were rehydrated with 0.1% FA and ∼600ng of sample was loaded onto a Evotip Pure™ Tip following the manufacturer’s protocol. The peptides were separated by the Xcalibur 15 SPD Extended gradient method (88min gradient), Evosep EV1106 column C18 15cmX150µm,1.9µm 30µm ID stainless emitter tip) using a Evosep One system (Evosep Biosystems, Odense, Denmark). The chromatography system was coupled online with an Orbitrap Fusion Tribrid mass spectrometer (Thermo Fisher Scientific, San Jose, CA) equipped with a Nanospray Flex NG source (Thermo Fisher Scientific). Solvents were A: 0.1% Formic acid/Water; B: 100% Acetonitrile, 0.1% Formic acid. The Orbitrap Fusion instrument parameters (Fusion Tune 4.2.4321 software) were as follows for orbitrap (OT-MS) orbitrap (OT-MS/MS) with HCD fragmentation: spray voltage 2050V, capillary temperature 300 ℃. The acquired survey MS1 scan range was 350-1550 m/z profile mode, resolution 60,000 FWHM@200m/z one microscan with Automatic inject time with the Siloxane mass 445.12002 used as lock mass for internal calibration. Data-dependent acquisition Orbitrap survey spectra were scheduled at least every 3 seconds, with the software determining “Top number most intense” ions for MS/MS acquisitions during this period. The automatic gain control (AGC) target value for FTMS was set to Standard 100% (400,000 counts) and automatic maximum fill time to ensure optimal sensitivity and cycle time. The most intense ions charge state 2-4 exceeding 50,000 counts were selected for HCD MSMS fragmentation in the ion routing multipole. Monoisotopic Precursor Selection (MIPS) most abundant peptide peak was enabled; Dynamic exclusion settings were repeat count: 2; repeat duration: 3 seconds; exclusion duration: 5 seconds with a 10ppm mass window. The data dependent (ddMS2) OT HCD scan used a quadrupole isolation window of 1.6 Da; Orbitrap detection at 30,000 resolution, auto mode normal m/z range, centroid detection, 1 microscan, auto maximum injection time, Custom AGC target (800%) or 400000 counts and stepped HCD collision energy of 30 and 35%.

#### Evosep One Gradient Details

The 15 SPD method has an 88-minute gradient and a cycle time of 93 minutes. The analytical column is equilibrated at 1500 nl/min. The gradient flow is 220 nl/min and increased to 1500 nl/min for washing. The method used the EV1106 Endurance column at ambient temperature (23 °C) with a 30µm I.D. stainless steel emitter tip.

#### Peptide identification

MS/MS datasets were analysed using FragPipe v23.1 and peptide identification was carried out by using a false discovery-based approach using a database of purified proteins and known contaminants^61,62^. MSFragger was used, and the precursor mass tolerance error was set to −20 to 20 ppm. The fragment mass tolerance was set at 20 ppm. Protein digestion was set as strict trypsin, allowing two missed cleavages, searching between lengths of 7 and 50 aa, with a mass range of 500 to 5000 Da, searching for methionine oxidation, N-terminal acetylation, and STY phosphorylation variable modifications.

### Lipid Vesicle Preparation

Two sets of lipid vesicles were prepared: 100% PI and ER-PM mimic vesicles (10% PI, 25% PS, 40% PE, 15% PC, 10% Cholesterol). Lipid vesicles were prepared by mixing the lipid dissolved in organic solvent. The solvent was evaporated in a stream of argon following which the lipid film was desiccated in a vacuum for 1 hour. The lipids were resuspended in lipid buffer (20 mM HEPES pH 7.5, 100 mM KCl and 0.5 mM EDTA) and the solution was sonicated for 15 minutes. The vesicles were subjected to ten freeze-thaw cycles and passed through an extruder (Avanti Research) with a 100 nm filter 21-31 times. The extruded vesicles were aliquoted and stored at −80°C.

### Kinase Assays

All lipid kinase activity assays employed the Transcreener ADP2 Fluorescence Intensity (FI) Assay (Bellbrook labs) which measures ADP production. For assays measuring differences in activity comparing ^WT^PI4KA to ^WT^PI4KA^pY1154,pY2090^, the final concentration of 100% PI vesicles was 0.5 mg/mL, and ATP was 100 μM ATP. Two μL of ^WT^PI4KA or ^WT^PI4KA^pY1154,pY2090^ at 2X final concentration (500 nM, 125 nM, 31.25 nM, 7.81 nM, 1.95 nM, 0.49 nM, 0.12 nM final in 1X) was mixed with 2 μL substrate solution containing ATP, and vesicles, and the reaction was allowed to proceed for 60 min at 20°C. The reaction was stopped with 4 μL of 2X stop and detect solution containing Stop and Detect buffer, 8 nM ADP Alexa Fluor 594 Tracer and 93.7 μg/mL ADP2 Antibody IRDye QC-1 and incubated for 60 minutes. The fluorescence intensity was measured using a SpectraMax M5 plate reader at excitation 590 nm and emission 620 nm. For assays measuring differences in activity between ^WT^PI4KA, ^WT^PI4KA^pY1154,pY2090^, ^Y2090F^PI4KA, ^Y2090F^PI4KA^pY1154^, ^Y1154F^PI4KA, and ^Y1154F^PI4KA^pY2090^, the final concentration of 100% PI vesicles was 0.5 mg/mL, and ATP was 100 μM ATP. Two μL of PI4KA complex solution at 2X final concentration (17.5 nM and 7 nM for ^WT^PI4KA, ^Y2090F^PI4KA, ^Y2090F^PI4KA^pY1154^, ^Y1154F^PI4KA, and 312.5 nM and 125 nM for ^WT^PI4KA^pY1154,pY2090^, and ^Y1154F^PI4KA^pY2090^) was mixed with 2 μL substrate solution containing ATP, and vesicles, and the reaction was allowed to proceed for 60 min at 20°C. The fluorescence intensity was measured using an Agilent BioTek synergy H1 plate reader at excitation 585 nm and emission 626 nm. For assays measuring differences in activity comparing ^WT^PI4KA to ^WT^PI4KA^pY1154,pY2090^, +/− 50 nM LactC2-EGFP-EFR3A on ER-PM mimic vesicles, the final concentration of ER-PM mimic vesicles was 0.5 mg/mL, and ATP was 100 μM ATP. Two μL of ^WT^PI4KA or ^WT^PI4KA^pY1154,pY2090^ at 2X final concentration (1000 nM, 250 nM, 62.5 nM, 15.63 nM, 3.91 nM, 0.98 nM, 0.24, and 0.06 nM final in 1X) was mixed with 2 μL substrate solution containing ATP, and ER-PM mimic vesicles +/− LactC2-EGFP-EFR3A (final 50 nM), and the reaction was allowed to proceed for 60 min at 20°C. The reaction was stopped with 4 μL of 2X stop and detect solution containing Stop and Detect buffer, 8 nM ADP Alexa Fluor 594 Tracer and 93.7 μg/mL ADP2 Antibody IRDye QC-1 and incubated for 60 minutes. The fluorescence intensity was measured using an Agilent BioTek synergy H1 plate reader at excitation 585 nm and emission 626 nm. Data was normalised against the appropriate 0–100% ADP window made using conditions containing either 100 μM ATP or ADP with kinase buffer. The percent ATP turnover was interpolated using a standard curve (0.1–100 μM ADP). Interpolated values were then used to calculate the specific activity of the enzymes.

### Cryo-EM

#### Cryo-EM sample preparation and data collection

C-Flat 2/1-3Cu-T-50 grids were glow-discharged for 25 s at 15 mA using a Pelco easiGlow glow discharger. 3 μL of ^WT^PI4KA^pY1154,pY2090^ were applied to the grids at 0.73 mg/mL and vitrified using a Vitrobot Mark IV (FEI) by blotting for 1.5 s at a blot force of −5 at 4°C and 100% humidity. Specimens were screened using a 200-kV Glacios transmission microscope (Thermo Fisher Scientific) equipped with a Falcon 3EC direct electron detector (DED). Datasets were collected using a 300-kV Titan Krios equipped with a Falcon 4i camera and the Selectris energy filter. A total of 23,312 super-resolution movies from two grids were collected using SerialEM with a total dose of 50 e^−^/Å^2^ over 693 frames at a physical pixel size of 0.77 Å per pixel, using a defocus range of −0.5 to −2.5 μm, at 165,000× magnification. See **Table S1.**

#### Cryo-EM image processing

All data processing was carried out using cryoSPARC v4.4.1, V4.5.1, and V4.5.2. Movies were uploaded with EER Upsampling set to 1. Patch motion correction was applied to all movies to align the frames. The contrast transfer function (CTF) of the resulting micrographs was estimated using the patch CTF estimation job with default settings. Final particle stack was manually curated to contain only micrographs with a CTF fit resolution less than or equal to 10.

To generate an initial model, 289,691 particles were picked from 2528 micrographs using blob picking with a minimum and maximum diameter of 200 and 300, respectively. Particles were inspected using the inspect picks job to remove particles that picked ice contamination and were then extracted with a box size of 600 for a total of 196,996 particles. The particles were subjected to 2D classification with 40 online EM iterations. A total of 11 2D classes (consisting of 55,582 particles) were selected and used to pick particles from the complete dataset.

To generate a cryo-EM map from the complete dataset, the above 2D templates were low pass filtered to 20 Å and used to template pick 22,976 micrographs with the particle diameter set to 400 Å. 3,347,819 particles were picked and following inspection of particles 2,208,530 particles were extracted with a box size of 560 pixels which were used for *ab initio* reconstruction and heterogeneous refinement using three classes and C1 symmetry. 855,469 particles from the best refinement class underwent subsequent *ab initio* reconstruction and heterogeneous refinement with three classes. A total of 348,672 from the highest quality refinement were subjected to further *ab initio* reconstruction with class similarity set to 0.8 to identify particle classes with discrete differences, followed by heterogeneous reconstruction. 242,295 particles from the highest quality refinement were subjected to homogeneous refinement with C1 symmetry. These particles were re-extracted from re-curated micrographs, with the resulting 239,639 particles undergoing an additional round of homogeneous refinement with C2 symmetry enforced, followed by C2 symmetry expansion of the particles. The symmetry-expanded particles were then used for a masked local refinement of the full map, yielding a reconstruction with an overall resolution of 2.98 Å based on the Fourier shell correlation (FSC) 0.143 criterion. The local resolution was determined with an FSC threshold of 0.143. The workflow to generate the final cryo-EM map is shown in **Fig S5**.

### Model building

The cryo-EM structure of PI4KA-TTC7B-FAM126A-EFR3A (PDB: 9BAX)^9^ was fit into the map using ChimeraX ^63^. EFR3A was removed from the model and underwent refinement in Phenix.real_space_refine ^64^ using realspace, rigid body, and adp refinement with tight secondary structure restraints, and manual building in COOT ^65^. Density of Y1154 allowed for modelling of a phosphate group at this residue. Refinements following the addition of a phosphate group on Y1154 were completed with occupancy of the phosphate group manually fixed to 50%. See **Table S1**.

### AlphaFold3

We modelled the pY2090 PI4KA kα12 helix using the AlphaFold server (https://www.alphafoldserver.com) ^66–68^ and input the sequences of PI4KA/TTC7B/FAM126. The resulting top ranked model had ptm and iptm scores of 0.8 and 0.78, respectively, and were consistent with cryo-EM data. The predicated aligned error (PAE) for the top-ranked output is shown in **Fig. S7**.

### Hydrogen-Deuterium Exchange Mass Spectrometry

#### HDX-MS Sample preparation

Individual HDX reactions comparing ^WT^PI4KA to ^WT^PI4KA^pY1154,pY2090^ were carried out in 10 µL reaction volumes containing 12 pmol of protein (1.2 µM final concentration of PI4KA complex). The exchange reactions were initiated by the addition of 8.2 µL of D_2_O buffer (20 mM Imidazole pH 7, 150 mM NaCl, 0.5 mM TCEP) to 1.8 µL of protein (final D_2_O concentration of 76.5 % [v/v]). Reactions proceeded for 3s, 30s, 300s and 3000s, at 20°C before being quenched with ice cold acidic quench buffer, resulting in a final concentration of 0.6M guanidine HCl and 0.9% formic acid post quench. All conditions and timepoints were created and run in independent triplicate. All samples were flash frozen immediately after quenching and stored at −80°C.

### HDX-MS Protein digestion and MS/MS data collection

Protein samples were rapidly thawed and injected onto an integrated fluidics system containing a HDx-3 PAL liquid handling robot and climate-controlled (2°C) chromatography system (Trajan), a Dionex Ultimate 3000 UHPLC system, as well as an Impact HD QTOF Mass spectrometer (Bruker) or Impact II QTOF Mass spectrometer (Bruker). The full details of the automated LC system are described in ^69^. The PI4KA complex were run over two immobilized pepsin columns (Affipro pepsin column, 69.3 μL, 2.1 mm × 20 mm; Affipro AP-PC-001) at 200 µL/min for 4 minutes at 2°C. The resulting peptides were collected and desalted on a C18 trap column (Acquity UPLC BEH C18 1.7mm column (2.1 mm x 5 mm); Waters 186004629). The trap was subsequently eluted in line with an ACQUITY 1.7 μm particle, 100 mm × 2.1 mm C18 UPLC column (Waters; 186003686), using a gradient of 3-35% B (Buffer A 0.1% formic acid; Buffer B 100% acetonitrile) over 11 minutes immediately followed by a gradient of 35-80% over 5 minutes. Mass spectrometry experiments acquired over a mass range from 150 to 2200 m/z using an electrospray ionization source operated at a temperature of 200°C and a spray voltage of 4.5 kV.

Experiments quantifying phosphorylation of Y1154 and/or Y2090 in the Lck dose response and in ^WT^PI4KA and ^WT^PI4KA^pY1154,pY2090^ used the same protein digestion and MS/MS data collection as above. Peptides from phosphorylation quantification ^Y2090F^PI4KA, ^Y2090F^PI4KA^pY1154^, ^Y1154F^PI4KA, ^Y1154F^PI4KA^pY2090^, AF647-^WT^PI4KA and AF647-^WT^PI4KA^pY1154,pY2090,^ as well as tracking increase in pY1154 intensity upon increasing concentrations of Lck, were eluted from the trap in line with an ACQUITY 1.7 µm particle, 2.1 mm × 100 mm C18 UPLC column (Waters; 186003686), using a gradient of 3-35% B (Buffer A 0.1% formic acid; Buffer B 100% acetonitrile) over 24 minutes immediately followed by a gradient of 35-80% over 2 minutes.

#### HDX-MS Peptide identification

Peptides were identified from non-deuterated samples using data-dependent acquisition following tandem MS/MS experiments (0.5s precursor scan from 150-2000 m/z; twelve 0.25s fragment scans from 150-2000 m/z). PI4KA complex MS/MS datasets were analysed using FragPipe v19.1 and peptide identification was carried out by using a false discovery-based approach using a database of purified proteins and known contaminants ^61,62^. MSFragger was used, and the precursor mass tolerance error was set to −20 to 20 ppm. The fragment mass tolerance was set at 20 ppm. Protein digestion was set as nonspecific, searching between lengths of 4 and 50 aa, with a mass range of 400 to 5000 Da.

#### HDX-MS Analysis of Peptide Centroids and Measurement of Deuterium Incorporation

HD-Examiner Software (Sierra Analytics) was used to automatically calculate the level of deuterium incorporation into each peptide. All peptides were manually inspected for correct charge state, correct retention time, appropriate selection of isotopic distribution, etc. Deuteration levels were calculated using the centroid of the experimental isotope clusters. Results are presented as relative levels of deuterium incorporation, and the only control for back exchange was the level of deuterium present in the buffer (76.5 %). Differences in exchange in a peptide were considered significant if they met all three of the following criteria: ≥5% change in exchange, ≥0.4 Da difference in exchange, and a p value <0.01 using a two tailed student t-test. The entire HDX-MS dataset with all the values and statistics are provided in the **source data**. Samples were only compared within a single experiment and were never compared to experiments completed at a different time with a different final D_2_O level. The data analysis statistics for all HDX-MS experiments are provided in the source data according to the guidelines of ^70^ (**Source Data**). All proteomics data generated in this study have been deposited to the ProteomeXchange Consortium via the PRIDE partner repository with the dataset identifiers PXD070483 and PXD083717 ^71^.

#### Bio-layer interferometry (BLI)

The BLI measurements were conducted using a Fortebio (Sartorius) K2 Octet using fiber optic biosensors according to the protocols of ^72^. Anti–penta-His biosensors were loaded using purified MBP-EFR3A (721-791), which had a 10x His tag on the N-terminus. The biosensor tips were preincubated in the BLI buffer (20 mM HEPES [pH 7.5], 150 mM NaCl, 0.01%, bovine serum albumin, and 0.002% Tween-20) for 10 min before experiments began. The sequence of steps in each assay was regeneration, custom, loading, baseline, association, and dissociation. Every experiment was done at 25 °C with shaking at 1000 rpm. Technical replicates were performed by using the same fiber tip and repeating the steps outlined previously. Regeneration was performed by exposing the tips to regeneration buffer (Glycine pH 1.5) for 5s and BLI buffer for 5s and repeating the exposure for 6 cycles. BLI buffer was used for the custom, baseline, and dissociation steps; these steps were performed in the same well for a given sample. To determine affinity constants, MBP-EFR3A (721-791) was diluted in BLI buffer to 50 nM and was loaded onto the anti penta-His biosensor tips. ^WT^PI4KA or ^WT^PI4KA^pY1154,pY2090^ were diluted in BLI buffer to 5 nM and added to the appropriate association wells. Non-specific association was controlled by with a no loading control (BLI buffer) and subtracting its responses from the responses measured with ^WT^PI4KA or ^WT^PI4KA^pY1154,pY2090^. No non-specific association of ^WT^PI4KA or ^WT^PI4KA^pY1154,pY2090^ with the no loading control was observed. The data were fit to a 1:1 model using the “full (assoc and dissoc) setting”. Reported kinetic binding constants (K_d_, k_ON_, and k_OFF_) values were calculated by taking the geometric mean of binding constants calculated for individual conditions and replicates.

### Statistics and Reproducibility

All experiments were carried out in triplicate, apart from cryo-EM structure determination and ^WT^PI4KA treatment with increasing amounts of Lck tracking gain of pY1154 peptide signal which were n=1. For phosphorylation quantification of ^WT^PI4KA treatment with SRC, one sample was removed from the analysis due to sample carryover. Where applicable, statistical analysis between conditions was performed using a two-tailed Student’s t-test.

## Data Availability

The EM data have been deposited in the EM data bank with accession EMD-73828 and the associated structural model has been deposited to the PDB with accession number PDB: 9Z5X (pdb_00009z5x). The MS proteomics data have been deposited to the ProteomeXchange Consortium via the PRIDE partner repository with the dataset identifiers PXD070483 and PXD083717 ^71^. All raw data in all figures are available in the source data excel file. All data needed to evaluate the conclusions in the paper are present in the paper and/or the Supplementary Materials.

## Code Availability

This study reports no original code.

## Supporting Information

This article contains supporting information.

## Acknowledgements

Grids were prepared and data collected at the High Resolution Macromolecular Electron Microscopy (HRMEM) facility at the University of British Columbia (https://cryoem.med.ubc.ca). We thank Claire Atkinson and Natalie Strynadka. HRMEM is funded by the Canadian Foundation for Innovation and the British Columbia Knowledge Development Fund. We thank Darryl Hardie from the Uvic-Genome BC proteomics centre for his involvement in running the Orbitrap Fusion samples. We thank Gerald Hammond and Tamas Balla for critical review of the manuscript before submission.

## Funding and Additional Information

J.E.B. is supported by the Canadian Institutes of Health Research (CIHR) (PJT-195808). A.L.S. is supported by a CIHR CGS-D scholarship. C.K.Y. is supported by CIHR (PJT-168907) and the Natural Sciences and Engineering Research Council of Canada (NGPIN-2024-04139). HRMEM is funded by the Canadian Foundation for Innovation and the British Columbia Knowledge Development Fund. S.D.H is supported by the National Science Foundation CAREER Award (MCB-2048060).

## Declaration of competing interest

The authors declare that they have no conflicts of interest with the contents of this article.

## CRediT authorship contribution statement

Alexandria L Shaw: Conceptualization, data curation, formal analysis, investigation, methodology, visualization, writing - original draft, writing - review and editing. Sophia Doerr: Data curation, formal analysis, investigation, methodology, writing - review and editing. Hunter G Nyvall: Formal Analysis, Investigation. Meredith L Jenkins: Formal Analysis, Investigation. Sushant Suresh: Investigation. Calvin K Yip: Funding acquisition, resources, supervision. Scott D Hansen: Data curation, formal analysis, funding acquisition, methodology, supervision, writing - review and editing. John E Burke: Conceptualization, funding acquisition, methodology, project administration, supervision, writing - original draft, writing - review and editing.

## Supplemental Figures and Tables

**Figure S1.**
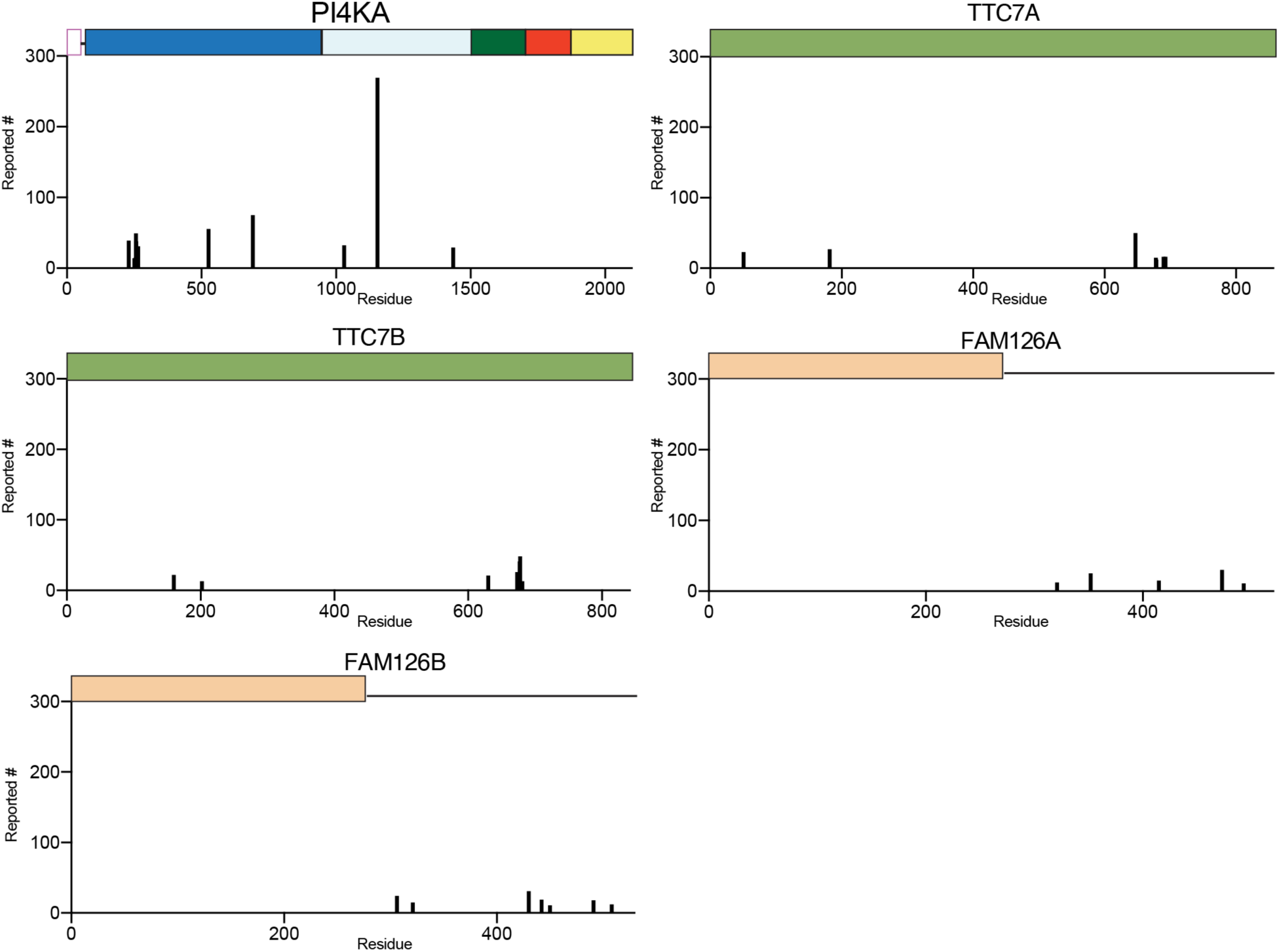
Reported phosphorylation sites for the PI4KA complex annotated in PhosphoSitePlus. Annotated phosphorylation sites in PI4KA, TTC7A/B, and FAM126A/B from PhosphoSitePlus_15_. Sites reported <5 times are not shown.

**Figure S2.**
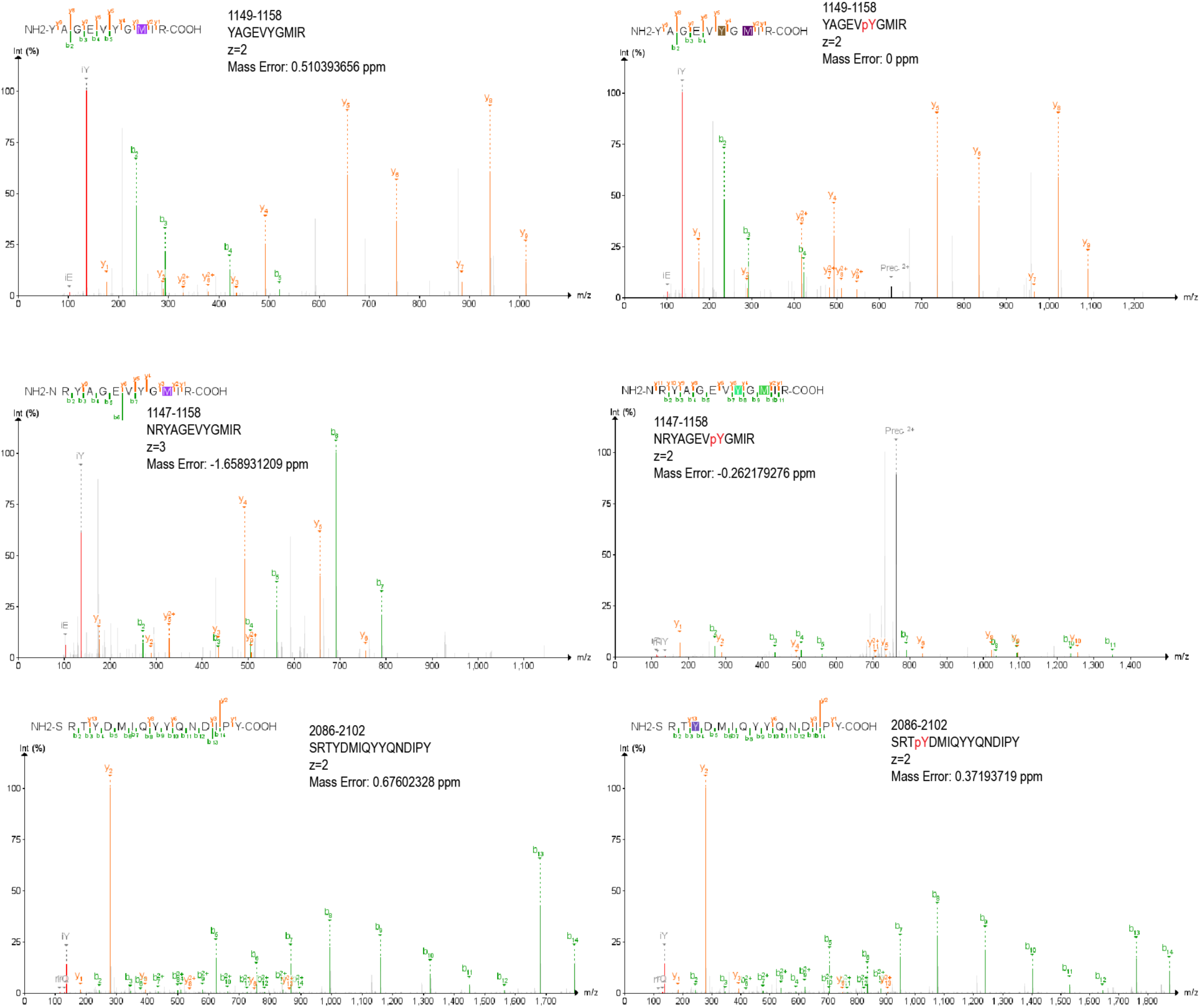
MS/MS spectra of peptides spanning Y1154 and Y2090 for the phosphorylated state. Identification of phosphorylated peptide species generated by tryptic digestion and analyzed by tandem mass spectrometry (MS/MS) using higher-energy collisional dissociation (HCD). The theoretical and experimentally observed precursor m/z values for each peptide are found in the source data.

**Figure S3.**
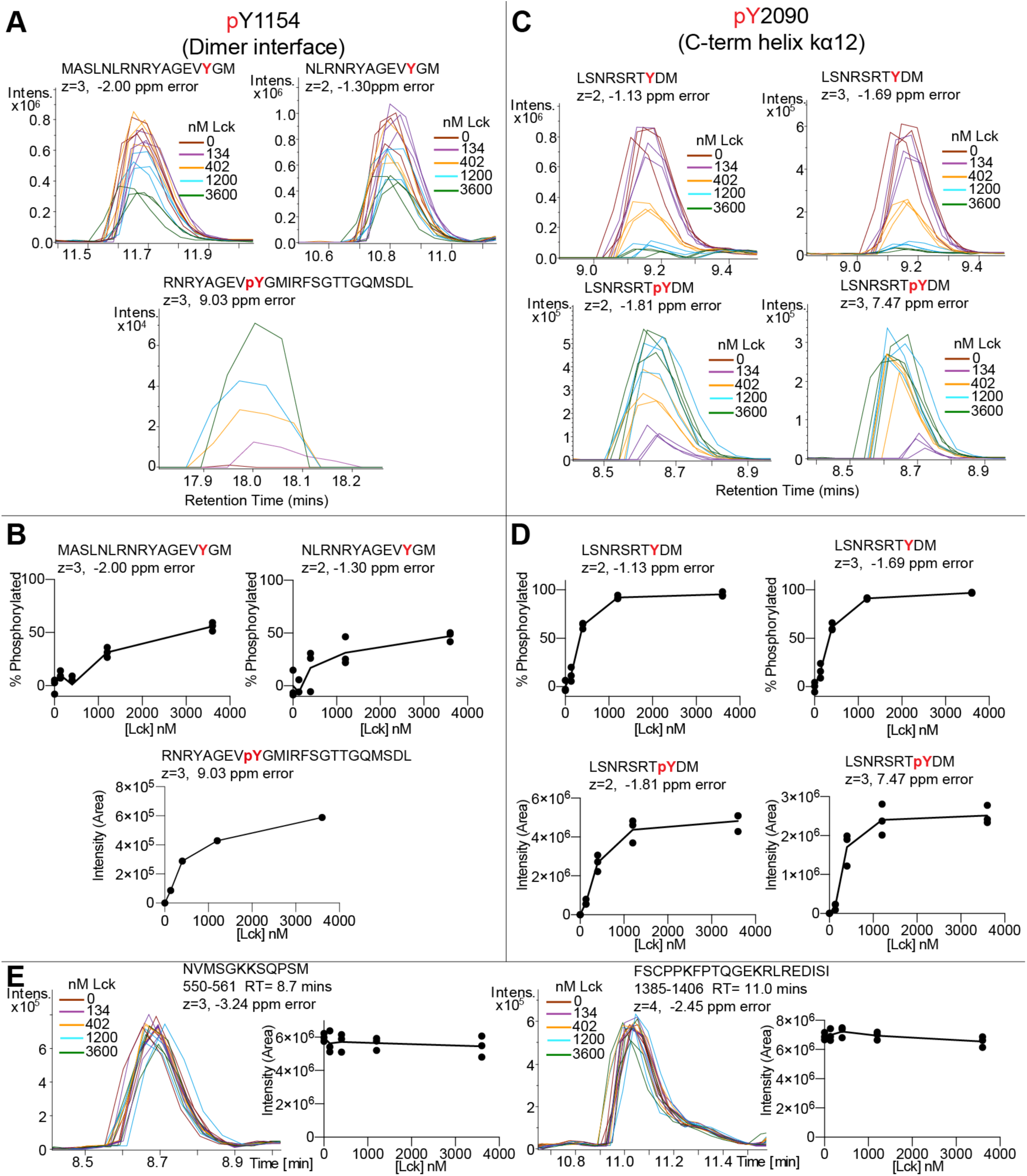
Loss of non-phosphorylated Y1154 and Y2090 PI4KA peptide intensity upon Lck treatment and gain of pY1154 and pY2090 PI4KA peptide intensity. Extracted ion chromatograms of two non-phosphorylated Y1154 (A) or Y2090 (C) containing peptides from samples with increasing concentrations of Lck (225-509) according to the legend, as well as the gain of a pY1154 and pY2090 peptide upon treatment with Lck (225-509). % Phosphorylation was calculated by tracking the loss of non-phosphorylated peptides. (B) (top) Y1154 was determined to be ∼50% phosphorylated across two unique peptides. An increase in a (B) (bottom) pY1154 containing peptide was also found and follows the same trend (n=1). (D) (top) Y2090 was determined to be >90% phosphorylated across two unique peptides with an increase in signal of corresponding pY2090 peptides (bottom). (E) Control PI4KA peptides that had similar retention times to the peptides in panels A-D are displayed to show that loss of Y1154 and Y2090 peptide signal is not a result of adding more protein to the system. (Independent replicates, all data points are shown, n=3 unless otherwise specified).

**Figure S4.**
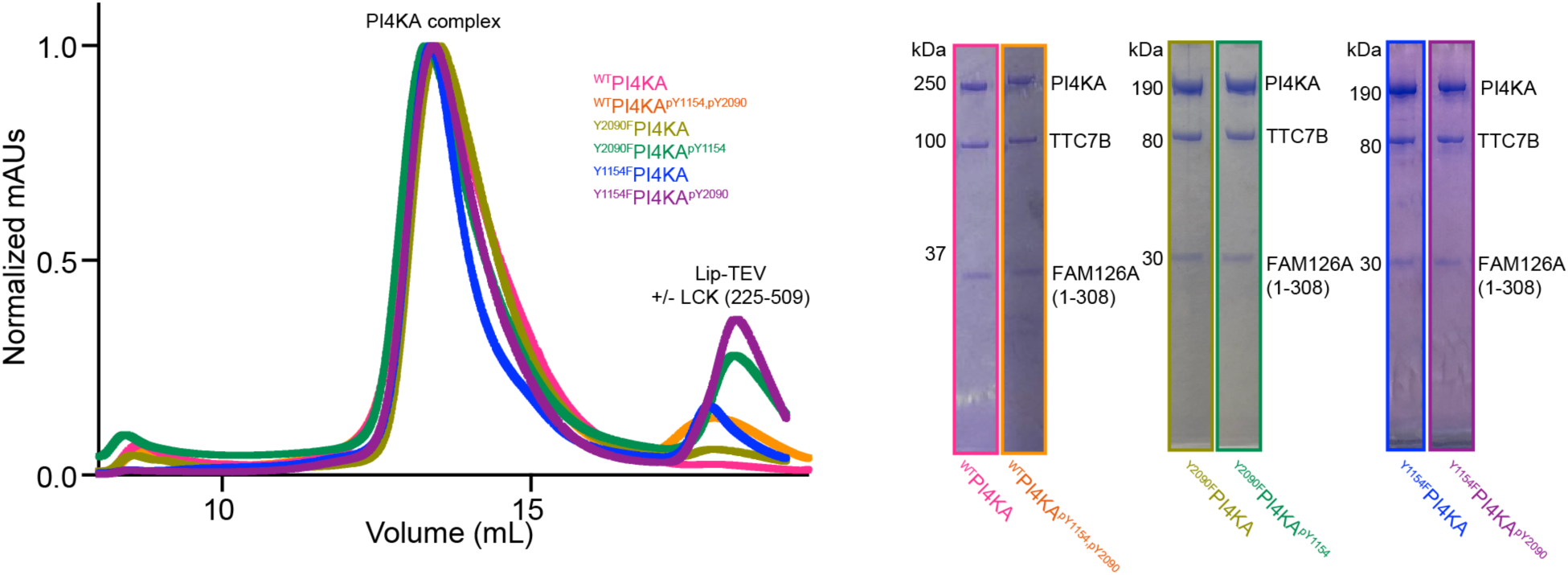
Protein purification of phosphorylated PI4KA complexes. Overlayed SEC traces (L) for all proteins in Figure 3D with their SDS-PAGE gels for the PI4KA complex peak(R).

**Figure S5:**
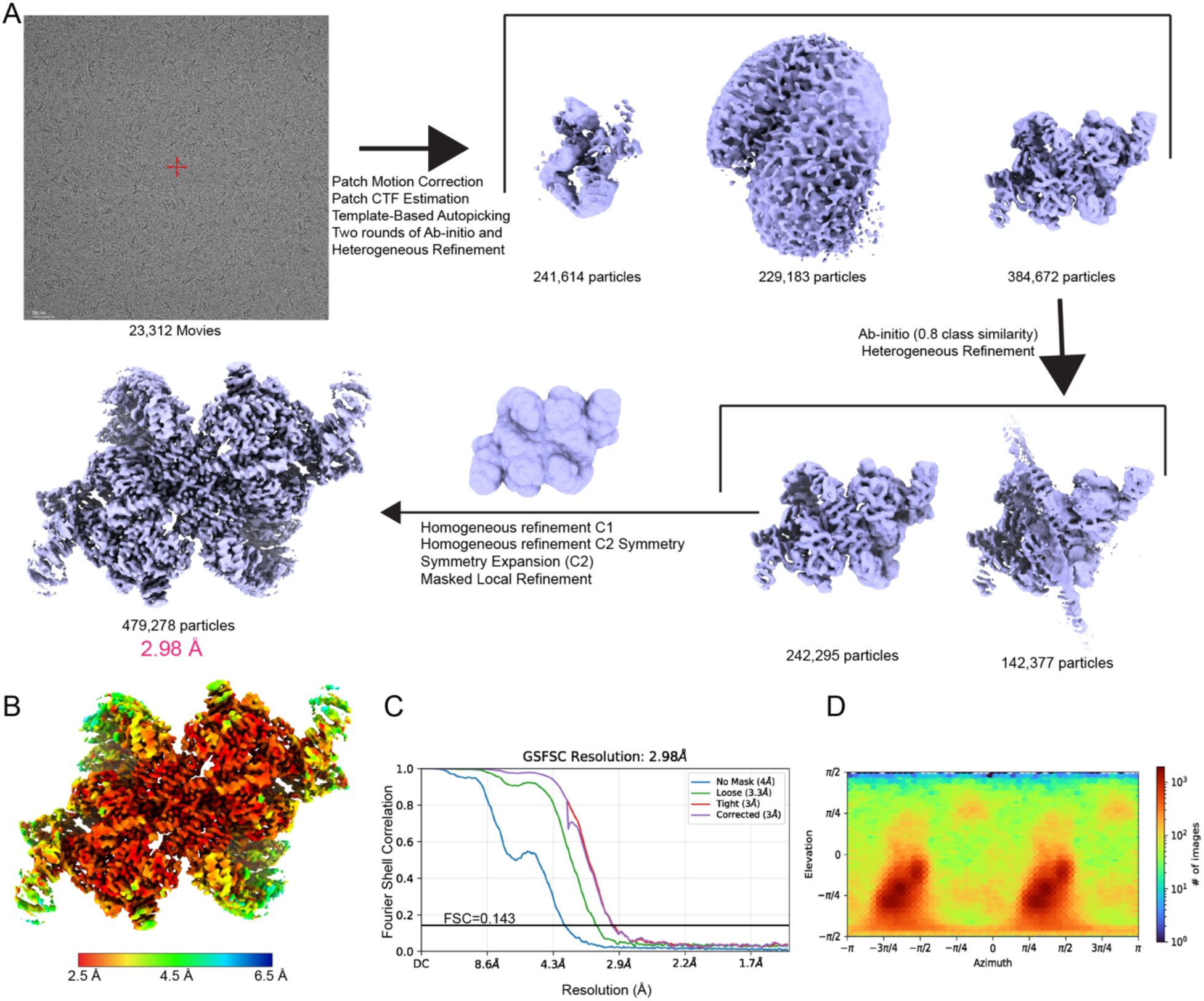
Cryo-EM data processing. A. Cryo-EM data processing workflow showing a representative micrograph from screening on the 200 kV Glacios, and the processing strategy used to generate a 3D reconstruction of ^WT^PI4KA^pY1154,pY2090^. B. Final map coloured according to local resolution estimated using cryoSPARC v4.5.2 (FSC=0.143). C. Gold standard Fourier shell correlation coefficient (FSC) curve after auto tightening by cryoSPARC for the final map. D. Viewing direction distribution plot of particles in the final cryo-EM reconstruction output by cryoSPARC v4.5.2.

**Figure S6:**
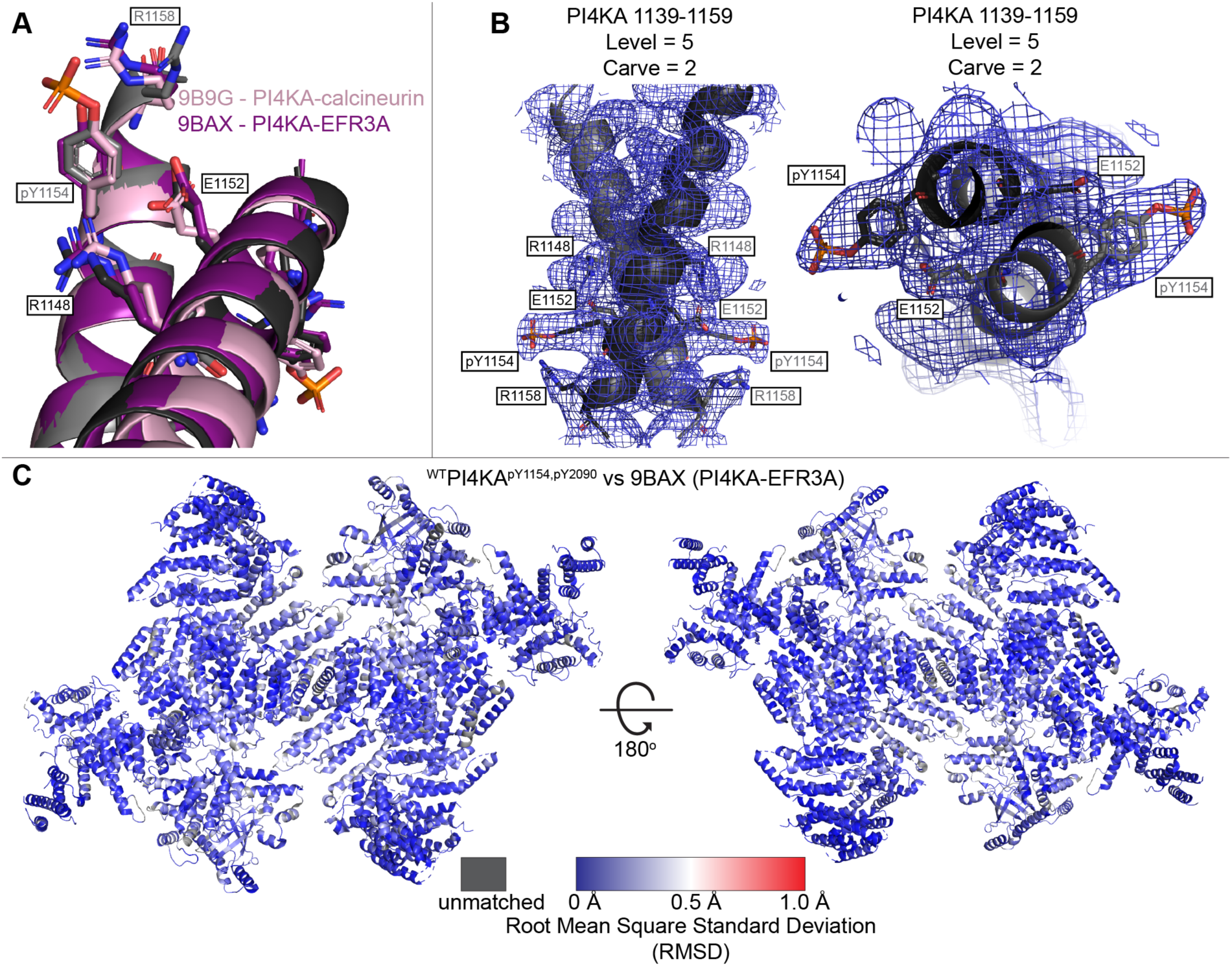
Model comparison and model density fit. A. Comparison of the ^WT^PI4KA^pY1154,pY2090^ structure from our study (grey) to PI4KA from PI4KA/TTC7B/FAM126A (1-308) with EFR3A (purple) (PDB:9BAX) and calcineurin (pink) (PDB:9B9G). Aligned on residues 1136-1159 of ^WT^PI4KA^pY1154,pY2090^. B. Electron density of PI4KA residues 1136-1159 with R1148, E1152, pY1154, and R1158 shown as sticks. C. RMSD comparison between ^WT^PI4KA^pY1154,pY2090^ structure and PI4KA/TTC7B/FAM126A (1-308) with EFR3A (PDB:9BAX) coloured according to the legend.

**Figure S7:**
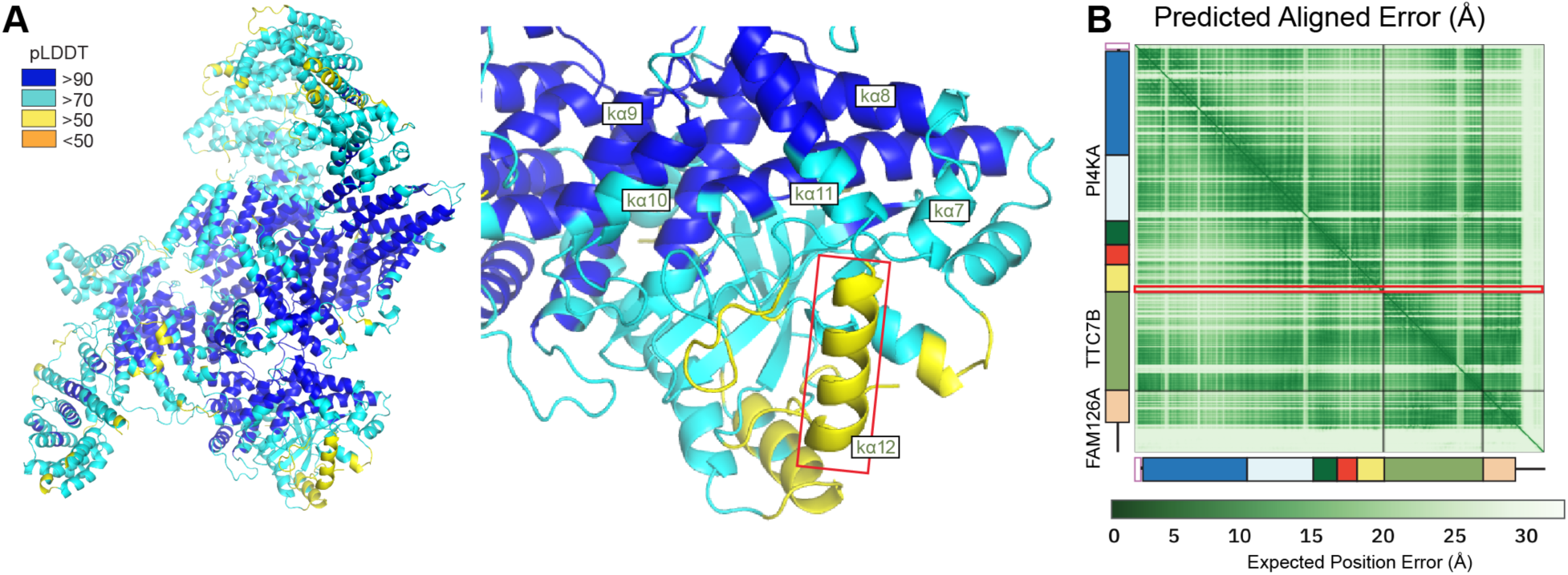
AlphaFold3 structure prediction of PI4KA pY2090 complex. A. AlphaFold3 prediction of PI4KA^pY2090^/TTC7B/FAM126A with the per-residue confidence metric predicted local-distance difference test shown as per the legend, with zoom-ins of the kinase domain’s regulatory motif. B. Predicted aligned error (PAE) of the AlphaFold3 prediction of PI4KA^pY2090^/TTC7B/FAM126A.

**Figure S8:**
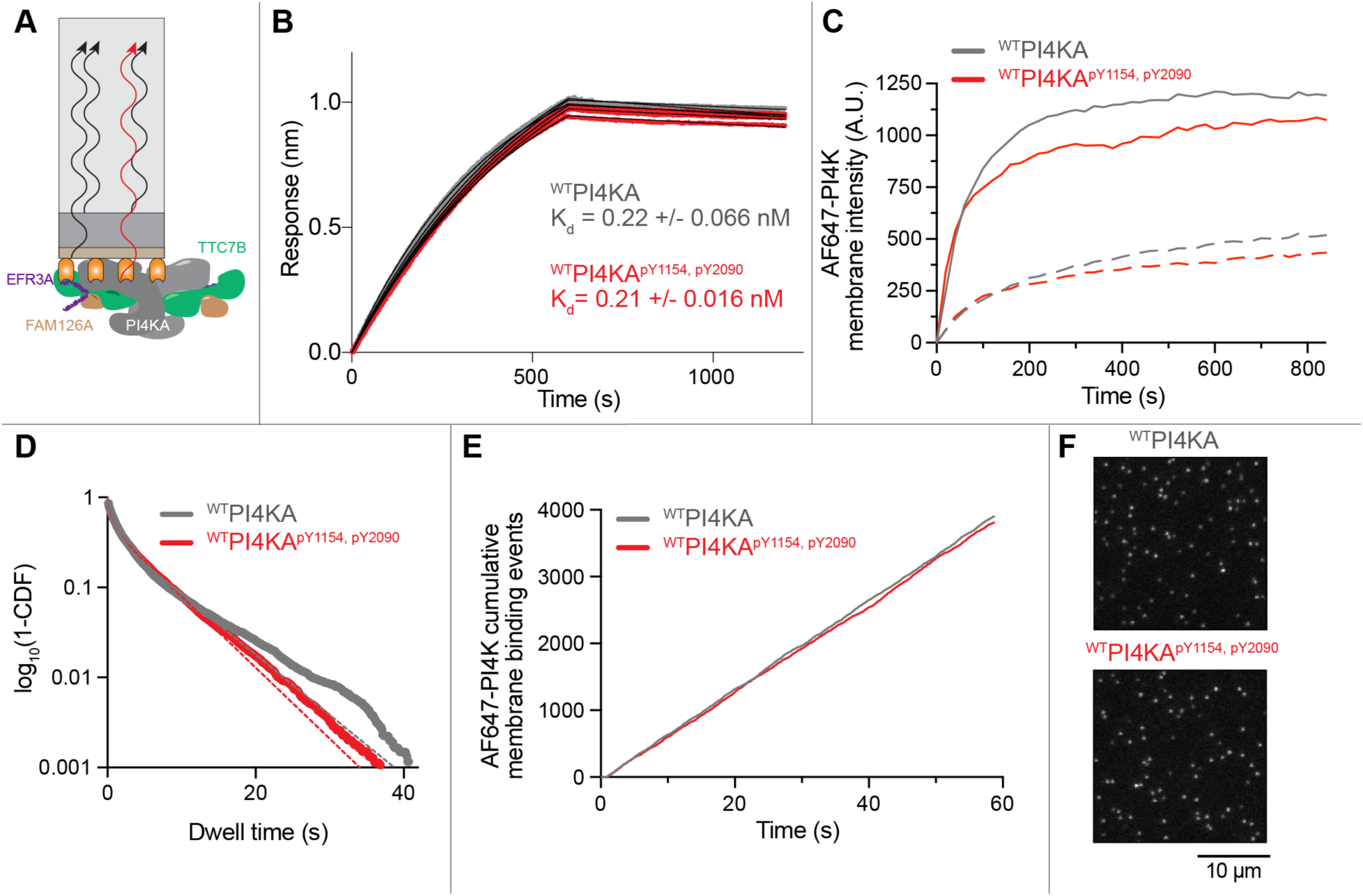
Phosphorylation of AF647-^WT^PI4KA does not perturb interactions with membrane tethered EFR3A. A. Cartoon of BLI experiment with EFR3A on the tip and ^WT^PI4KA or ^WT^PI4KA^pY1154,pY2090^ in solution. B. Raw BLI traces of EFR3A binding to ^WT^PI4KA or ^WT^PI4KA^pY1154,pY2090^ (5 nM). K_d_ values were calculated using a 1:1 binding model (N = 3 technical replicates). C. Bulk membrane absorption kinetics of ^WT^PI4KA and ^WT^PI4KA^pY1154,pY2090^ at 2 nM (solid lines) and 0.2 nM (dashed lines). D. Representative single molecule dwell time distributions measured in the presence of 200 pM AF647-PI4KA, +/− phosphorylation, on SLBs containing membrane-tethered EFR3A. Data plotted as log_10_(1-cumulative distribution frequency) and fit with a double exponential (dashed lines) yielded the following time constants: ^WT^PI4KA (τ_1_= 0.85 ± 0.06 s, τ_2_= 6.36 ± 0.28 s, α = 0.6, n = 14754 molecules) and ^WT^PI4KA^pY1154,pY2090^ (τ_1_= 0.63 ± 0.02 s, τ_2_= 5.55 ± 0.23 s, α = 0.5, n = 12120 molecules). Alpha (α) represents the fraction of molecules with a characteristic dwell time of τ_1_. Errors equal SD (N = 3 technical replicates). E. Phosphorylation of AF647-^WT^PI4KA does not modulate the single molecule membrane binding frequency. Representative cumulative membrane binding events were measured on a 3000 µm_2_ membrane surface coated with EFR3A. Data collect by single molecule TIRF microscopy in the presence of 200 pM AF647-^WT^PI4KA, yielding the following binding frequencies: 4.41 ± 0.2 AF647-^WT^PI4KA events • pM_-1_ • µm_-2_ • s_-1_ and 4.43 ± 0.2 AF647-^WT^PI4KA ^pY1154,pY2090^ events • pM_-1_ • µm_-2_ • s_-1_. Errors equal SD (N = 3 technical replicates). F. Representative TIRF-M images showing the localisation of the 200 pM AF647-^WT^PI4KA, +/− phosphorylation, on SLBs containing membrane-tethered EFR3A. C-F. Membrane composition: 86% DOPC, 10% PS, 2% PI, 2% MCC-PE (SpyCatcher conjugated).

**Table S1.**
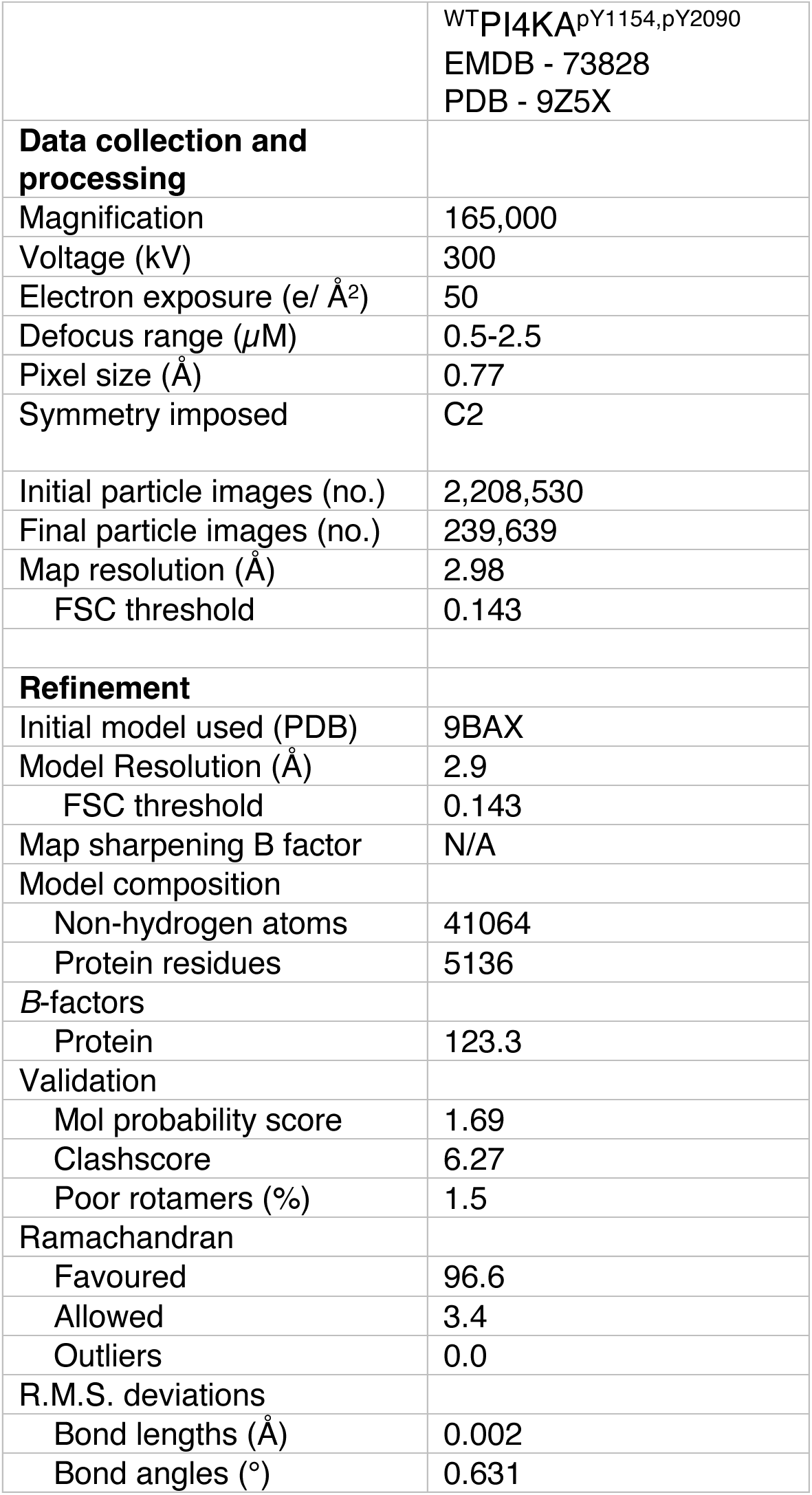
Cryo-EM data collection, refinement, and validation statistics.

**Table S2:**
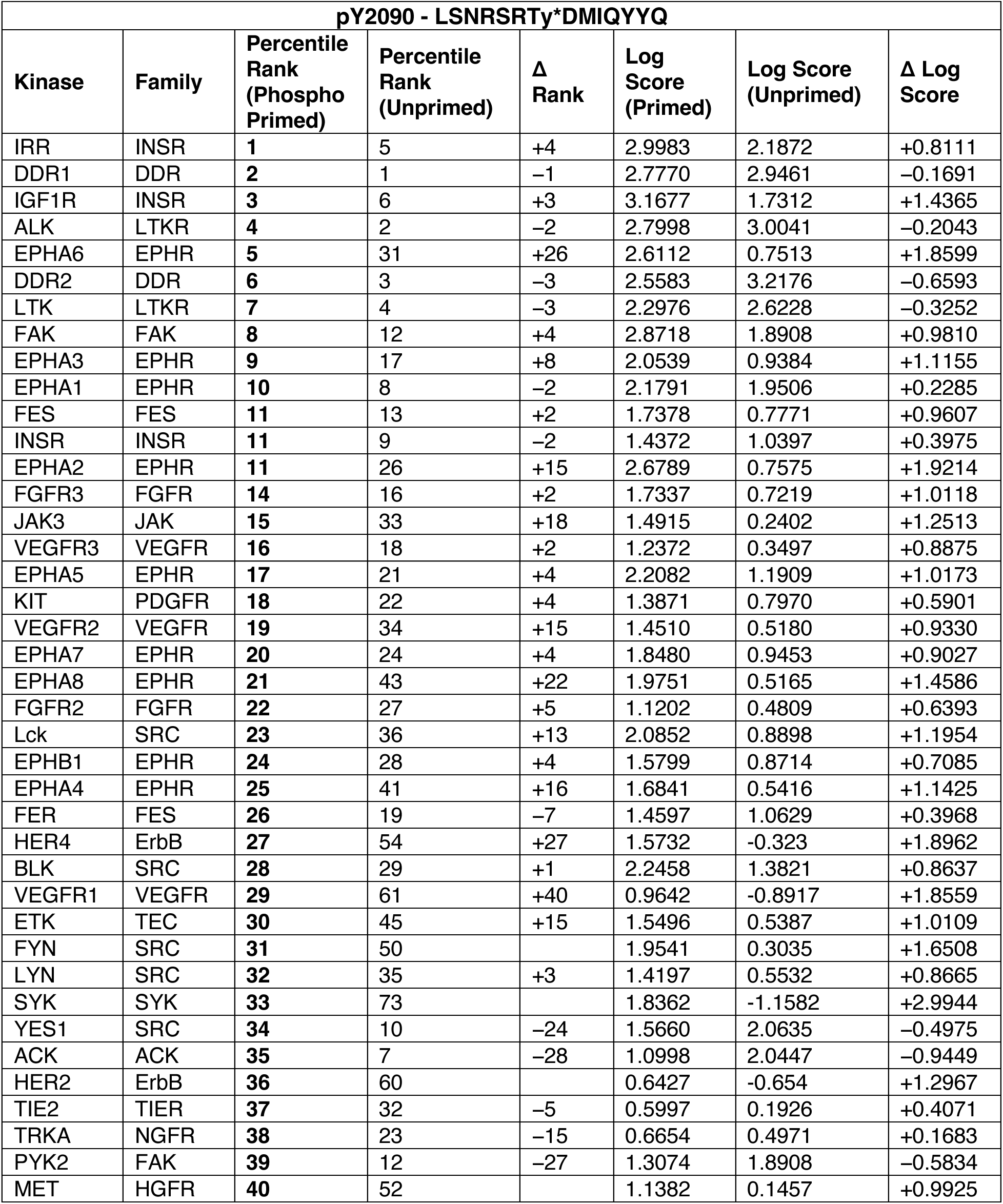
pY2090 predicted kinases enhanced with priming of T2089.

| pY2090 - LSNRSRTy*DMIQYYQ |  |  |  |  |  |  |  |
| --- | --- | --- | --- | --- | --- | --- | --- |
| Kinase | Family | Percentile Rank (Phospho Primed) | Percentile Rank (Unprimed) | Δ Rank | Log Score (Primed) | Log Score (Unprimed) | Δ Log Score |
| IRR | INSR | 1 | 5 | +4 | 2.9983 | 2.1872 | +0.8111 |
| DDR1 | DDR | 2 | 1 | -1 | 2.7770 | 2.9461 | -0.1691 |
| IGF1R | INSR | 3 | 6 | +3 | 3.1677 | 1.7312 | +1.4365 |
| ALK | LTKR | 4 | 2 | -2 | 2.7998 | 3.0041 | -0.2043 |
| EPHA6 | EPHR | 5 | 31 | +26 | 2.6112 | 0.7513 | +1.8599 |
| DDR2 | DDR | 6 | 3 | -3 | 2.5583 | 3.2176 | -0.6593 |
| LTK | LTKR | 7 | 4 | -3 | 2.2976 | 2.6228 | -0.3252 |
| FAK | FAK | 8 | 12 | +4 | 2.8718 | 1.8908 | +0.9810 |
| EPHA3 | EPHR | 9 | 17 | +8 | 2.0539 | 0.9384 | +1.1155 |
| EPHA1 | EPHR | 10 | 8 | -2 | 2.1791 | 1.9506 | +0.2285 |
| FES | FES | 11 | 13 | +2 | 1.7378 | 0.7771 | +0.9607 |
| INSR | INSR | 11 | 9 | -2 | 1.4372 | 1.0397 | +0.3975 |
| EPHA2 | EPHR | 11 | 26 | +15 | 2.6789 | 0.7575 | +1.9214 |
| FGFR3 | FGFR | 14 | 16 | +2 | 1.7337 | 0.7219 | +1.0118 |
| JAK3 | JAK | 15 | 33 | +18 | 1.4915 | 0.2402 | +1.2513 |
| VEGFR3 | VEGFR | 16 | 18 | +2 | 1.2372 | 0.3497 | +0.8875 |
| EPHA5 | EPHR | 17 | 21 | +4 | 2.2082 | 1.1909 | +1.0173 |
| KIT | PDGFR | 18 | 22 | +4 | 1.3871 | 0.7970 | +0.5901 |
| VEGFR2 | VEGFR | 19 | 34 | +15 | 1.4510 | 0.5180 | +0.9330 |
| EPHA7 | EPHR | 20 | 24 | +4 | 1.8480 | 0.9453 | +0.9027 |
| EPHA8 | EPHR | 21 | 43 | +22 | 1.9751 | 0.5165 | +1.4586 |
| FGFR2 | FGFR | 22 | 27 | +5 | 1.1202 | 0.4809 | +0.6393 |
| Lck | SRC | 23 | 36 | +13 | 2.0852 | 0.8898 | +1.1954 |
| EPHB1 | EPHR | 24 | 28 | +4 | 1.5799 | 0.8714 | +0.7085 |
| EPHA4 | EPHR | 25 | 41 | +16 | 1.6841 | 0.5416 | +1.1425 |
| FER | FES | 26 | 19 | -7 | 1.4597 | 1.0629 | +0.3968 |
| HER4 | ErbB | 27 | 54 | +27 | 1.5732 | -0.323 | +1.8962 |
| BLK | SRC | 28 | 29 | +1 | 2.2458 | 1.3821 | +0.8637 |
| VEGFR1 | VEGFR | 29 | 61 | +40 | 0.9642 | -0.8917 | +1.8559 |
| ETK | TEC | 30 | 45 | +15 | 1.5496 | 0.5387 | +1.0109 |
| FYN | SRC | 31 | 50 |  | 1.9541 | 0.3035 | +1.6508 |
| LYN | SRC | 32 | 35 | +3 | 1.4197 | 0.5532 | +0.8665 |
| SYK | SYK | 33 | 73 |  | 1.8362 | -1.1582 | +2.9944 |
| YES1 | SRC | 34 | 10 | -24 | 1.5660 | 2.0635 | -0.4975 |
| ACK | ACK | 35 | 7 | -28 | 1.0998 | 2.0447 | -0.9449 |
| HER2 | ErbB | 36 | 60 |  | 0.6427 | -0.654 | +1.2967 |
| TIE2 | TIER | 37 | 32 | -5 | 0.5997 | 0.1926 | +0.4071 |
| TRKA | NGFR | 38 | 23 | -15 | 0.6654 | 0.4971 | +0.1683 |
| PYK2 | FAK | 39 | 12 | -27 | 1.3074 | 1.8908 | -0.5834 |
| MET | HGFR | 40 | 52 |  | 1.1382 | 0.1457 | +0.9925 |

**Table S3.**
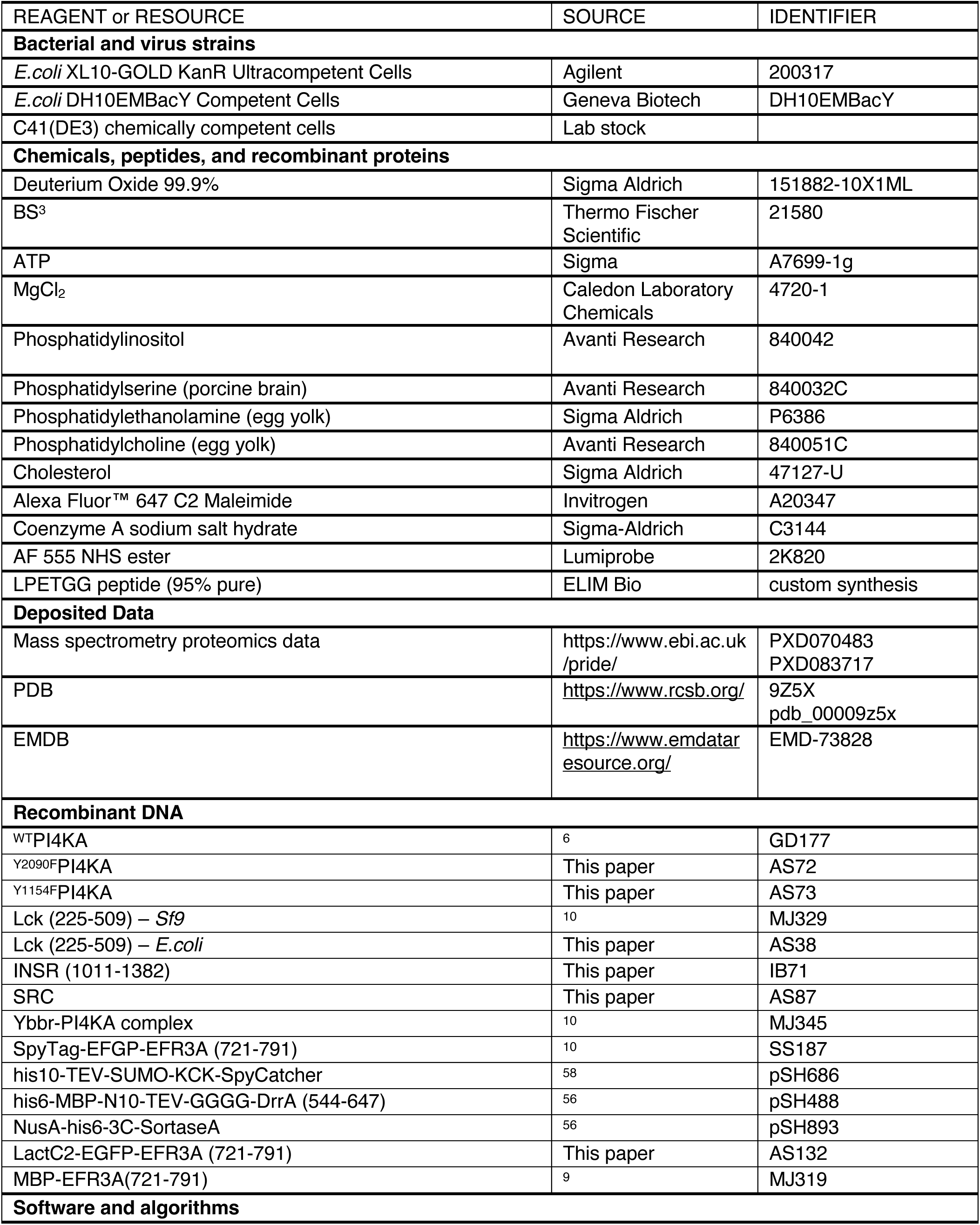
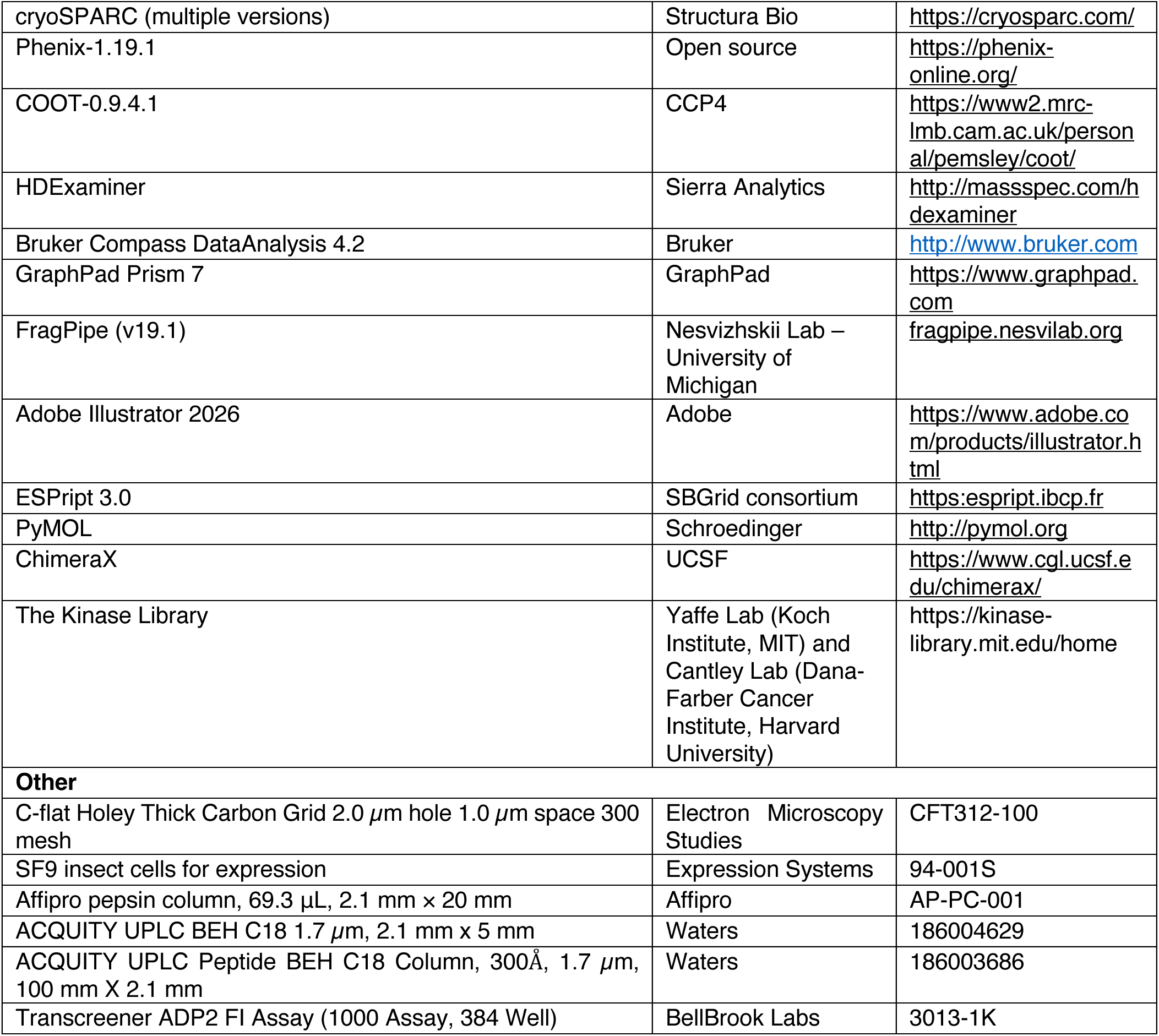
Key Reagents/Resources.

## Notes

### Competing Interest Statement

The authors have declared no competing interest.

### Summary of Updates

Additional experimentation for quantification and identification of the Y2090 phosphosite

## References

1. Balla, T. Phosphatidylinositol 4-phosphate; A minor lipid with multiple personalities. Biochim Biophys Acta Mol Cell Biol Lipids 1870, 159615 (2025).

2. Chung, J. et al. PI4P/phosphatidylserine countertransport at ORP5- and ORP8-mediated ER-plasma membrane contacts. Science 349, 428–432 (2015).

3. Balla, T. Phosphoinositides: tiny lipids with giant impact on cell regulation. Physiol Rev 93, 1019–1137 (2013).

4. Lees, J. A. et al. Architecture of the human PI4KIIIα lipid kinase complex. Proceedings of the National Academy of Sciences of the United States of America 114, 13720–13725 (2017).

5. Baskin, J. M. et al. The leukodystrophy protein FAM126A (hyccin) regulates PtdIns(4)P synthesis at the plasma membrane. Nat Cell Biol 18, 132–138 (2016).

6. Dornan, G. L. et al. Probing the Architecture, Dynamics, and Inhibition of the PI4KIIIα/TTC7/FAM126 Complex. Journal of Molecular Biology 430, 3129–3142 (2018).

7. Bojjireddy, N., Guzman-Hernandez, M. L., Reinhard, N. R., Jovic, M. & Balla, T. EFR3s are palmitoylated plasma membrane proteins that control responsiveness to G-protein-coupled receptors. J Cell Sci 128, 118–128 (2015).

8. Baird, D., Stefan, C., Audhya, A., Weys, S. & Emr, S. D. Assembly of the PtdIns 4-kinase Stt4 complex at the plasma membrane requires Ypp1 and Efr3. J Cell Biol 183, 1061–1074 (2008).

9. Suresh, S. et al. Molecular basis for plasma membrane recruitment of PI4KA by EFR3. Sci Adv 10, eadp6660 (2024).

10. Suresh, S. et al. Development of an inhibitory TTC7B selective nanobody that blocks EFR3 recruitment of PI4KA. J Biol Chem 301, 110886 (2025).

11. Chung, J., Nakatsu, F., Baskin, J. M. & De Camilli, P. Plasticity of PI4KIIIα interactions at the plasma membrane. EMBO Rep. 16, 312–320 (2015).

12. Ulengin-Talkish, I. et al. Palmitoylation targets the calcineurin phosphatase to the phosphatidylinositol 4-kinase complex at the plasma membrane. Nat Commun 12, 6064 (2021).

13. Shaw, A. L. et al. Structure of calcineurin bound to PI4KA reveals dual interface in both PI4KA and FAM126A. Structure S0969-2126(24)00324–1 (2024) doi:10.1016/j.str.2024.08.007.

14. Xŭ, X. J., Tong, C. S. & Wu, M. Distinct impact of PI(4)P flux on PI(4,5)P2 steady states and oscillations. Proc Natl Acad Sci U S A 123, e2518354123 (2026).

15. Hornbeck, P. V. et al. PhosphoSitePlus, 2014: mutations, PTMs and recalibrations. Nucleic Acids Res 43, D512–520 (2015).

16. Burke, J. E. Structural Basis for Regulation of Phosphoinositide Kinases and Their Involvement in Human Disease. Mol. Cell 71, 653–673 (2018).

17. Miller, S. et al. Shaping development of autophagy inhibitors with the structure of the lipid kinase Vps34. Science 327, 1638–1642 (2010).

18. Burke, J. E. et al. Structures of PI4KIIIβ complexes show simultaneous recruitment of Rab11 and its effectors. Science 344, 1035–1038 (2014).

19. Shaw, A. L., Barlow-Busch, I. & Burke, J. E. Molecular basis for regulation of the class I phosphoinositide 3-kinases (PI3Ks), and their targeting in human disease. Biochim Biophys Acta Mol Cell Biol Lipids 1870, 159689 (2025).

20. Torosyan, H. et al. Structures of the PI3Kα/KRas complex on lipid bilayers reveal molecular mechanisms of PI3Kα activation. Mol Cell 86, 2858–2870.e6 (2026).

21. Czupalla, C. et al. Identification and characterization of the autophosphorylation sites of phosphoinositide 3-kinase isoforms beta and gamma. J Biol Chem 278, 11536–11545 (2003).

22. Vanhaesebroeck, B. et al. Autophosphorylation of p110delta phosphoinositide 3-kinase: a new paradigm for the regulation of lipid kinases in vitro and in vivo. EMBO J. 18, 1292–1302 (1999).

23. Lin, T.-Y. et al. Epinephrine inhibits PI3Kα via the Hippo kinases. Cell Rep 42, 113535 (2023).

24. Barlow-Busch, I., Shaw, A. L. & Burke, J. E. PI4KA and PIKfyve: Essential phosphoinositide signaling enzymes involved in myriad human diseases. Current Opinion in Cell Biology 83, 102207 (2023).

25. Berger, K. L., Kelly, S. M., Jordan, T. X., Tartell, M. A. & Randall, G. Hepatitis C virus stimulates the phosphatidylinositol 4-kinase III alpha-dependent phosphatidylinositol 4-phosphate production that is essential for its replication. J. Virol. 85, 8870–8883 (2011).

26. Bojjireddy, N. et al. Pharmacological and genetic targeting of the PI4KA enzyme reveals its important role in maintaining plasma membrane phosphatidylinositol 4-phosphate and phosphatidylinositol 4,5-bisphosphate levels. J. Biol. Chem. 289, 6120–6132 (2014).

27. Adhikari, H. et al. Oncogenic KRAS is dependent upon an EFR3A-PI4KA signaling axis for potent tumorigenic activity. Nat Commun 12, 5248 (2021).

28. Kattan, W. E. et al. Components of the phosphatidylserine endoplasmic reticulum to plasma membrane transport mechanism as targets for KRAS inhibition in pancreatic cancer. Proc Natl Acad Sci USA 118, e2114126118 (2021).

29. Yaron-Barir, T. M. et al. The intrinsic substrate specificity of the human tyrosine kinome. Nature 629, 1174–1181 (2024).

30. Shah, N. H., Löbel, M., Weiss, A. & Kuriyan, J. Fine-tuning of substrate preferences of the Src-family kinase Lck revealed through a high-throughput specificity screen. Elife 7, e35190 (2018).

31. Prasad, K. V. et al. Phosphatidylinositol (PI) 3-kinase and PI 4-kinase binding to the CD4-p56lck complex: the p56lck SH3 domain binds to PI 3-kinase but not PI 4-kinase. Mol Cell Biol 13, 7708–7717 (1993).

32. Zhang, Z. et al. Quantitative phosphoproteomics reveal cellular responses from caffeine, coumarin and quercetin in treated HepG2 cells. Toxicol Appl Pharmacol 449, 116110 (2022).

33. Lin, S. et al. EPSD: a well-annotated data resource of protein phosphorylation sites in eukaryotes. Brief Bioinform 22, 298–307 (2021).

34. Zhang, X. et al. Alterations in the Global Proteome and Phosphoproteome in Third Generation EGFR TKI Resistance Reveal Drug Targets to Circumvent Resistance. Cancer Res 81, 3051–3066 (2021).

35. Kitata, R. B. et al. A data-independent acquisition-based global phosphoproteomics system enables deep profiling. Nat Commun 12, 2539 (2021).

36. Zhang, Z. et al. Quantitative phosphoproteomics reveal cellular responses from caffeine, coumarin and quercetin in treated HepG2 cells. Toxicol Appl Pharmacol 449, 116110 (2022).

37. Johnson, J. L. et al. An atlas of substrate specificities for the human serine/threonine kinome. Nature 613, 759–766 (2023).

38. Woessner, N. M., Uleri, V., Stepanek, O. & Minguet, S. The TCR and LCK: foundations for T-cell activation and therapeutic innovation. Front Immunol 16, 1737013 (2025).

39. Luo, S., Du, S., Tao, M., Cao, J. & Cheng, P. Insights on hematopoietic cell kinase: An oncogenic player in human cancer. Biomed Pharmacother 160, 114339 (2023).

40. Pelaz, S. G. & Tabernero, A. Src: coordinating metabolism in cancer. Oncogene 41, 4917–4928 (2022).

41. Yeatman, T. J. A renaissance for SRC. Nat Rev Cancer 4, 470–480 (2004).

42. Trono, P., Ottavi, F. & Rosano’, L. Novel insights into the role of Discoidin domain receptor 2 (DDR2) in cancer progression: a new avenue of therapeutic intervention. Matrix Biol 125, 31–39 (2024).

43. Su, H. & Karin, M. Multifaceted collagen-DDR1 signaling in cancer. Trends Cell Biol 34, 406–415 (2024).

44. Voena, C., Ambrogio, C., Iannelli, F. & Chiarle, R. ALK in cancer: from function to therapeutic targeting. Nat Rev Cancer 25, 359–378 (2025).

45. Yeung, T. et al. Membrane phosphatidylserine regulates surface charge and protein localization. Science 319, 210–213 (2008).

46. Nakatsu, F. et al. PtdIns4P synthesis by PI4KIIIα at the plasma membrane and its impact on plasma membrane identity. J. Cell Biol. 199, 1003–1016 (2012).

47. Hammond, G. R. V. et al. PI4P and PI(4,5)P2 are essential but independent lipid determinants of membrane identity. Science 337, 727–730 (2012).

48. Hammond, G. R. V., Machner, M. P. & Balla, T. A novel probe for phosphatidylinositol 4-phosphate reveals multiple pools beyond the Golgi. J. Cell Biol. 205, 113–126 (2014).

49. Fowler, M. L. et al. Using hydrogen deuterium exchange mass spectrometry to engineer optimized constructs for crystallization of protein complexes: Case study of PI4KIIIβ with Rab11. Protein Sci. 25, 826–839 (2016).

50. Zhang, X. et al. Structure of Lipid Kinase p110b/p85b Elucidatesan Unusual SH2-Domain-Mediated Inhibitory Mechanism. Mol. Cell 41, 567–578 (2011).

51. Hon, W. C., Berndt, A. & Williams, R. L. Regulation of lipid binding underlies the activation mechanism of class IA PI3-kinases. Oncogene 31, 3655–3666 (2012).

52. Cook, A. S. I. et al. Structural pathway for PI3-kinase regulation by VPS15 in autophagy. Science 388, eadl3787 (2025).

53. Kamacioglu, A., Tuncbag, N. & Ozlu, N. Structural analysis of mammalian protein phosphorylation at a proteome level. Structure 29, 1219–1229.e3 (2021).

54. Gouet, P., Xavier, R. & Courcelle, E. ESPript/ENDscript: extracting and rendering sequence and 3D information from atomic structures of proteins. Nucleic Acids Research 31, 3320–3323 (2003).

55. Weissmann, F. et al. BiGBac enables rapid gene assembly for the expression of large multisubunit protein complexes. Proceedings of the National Academy of Sciences of the United States of America 113, E2564–E2569 (2016).

56. Hansen, S. D. et al. Stochastic geometry sensing and polarization in a lipid kinase– phosphatase competitive reaction. Proc. Natl. Acad. Sci. U.S.A. 116, 15013–15022 (2019).

57. Hansen, S. D., Lee, A. A., Duewell, B. R. & Groves, J. T. Membrane-mediated dimerization potentiates PIP5K lipid kinase activity. Elife 11, e73747 (2022).

58. Duewell, B. R., Faris, K. A. & Hansen, S. D. Molecular basis of product recognition during PIP5K-mediated production of PI(4,5)P2 with positive feedback. J Biol Chem 300, 107631 (2024).

59. Yin, J., Lin, A. J., Golan, D. E. & Walsh, C. T. Site-specific protein labeling by Sfp phosphopantetheinyl transferase. Nat Protoc 1, 280–285 (2006).

60. Jaqaman, K. et al. Robust single-particle tracking in live-cell time-lapse sequences. Nat Methods 5, 695–702 (2008).

61. Dobbs, J. M., Jenkins, M. L. & Burke, J. E. Escherichia coli and Sf9 Contaminant Databases to Increase Efficiency of Tandem Mass Spectrometry Peptide Identification in Structural Mass Spectrometry Experiments. J Am Soc Mass Spectrom 31, 2202–2209 (2020).

62. Kong, A. T., Leprevost, F. V., Avtonomov, D. M., Mellacheruvu, D. & Nesvizhskii, A. I. MSFragger: ultrafast and comprehensive peptide identification in mass spectrometry-based proteomics. Nat Methods 14, 513–520 (2017).

63. Pettersen, E. F. et al. UCSF CHIMERAX : Structure visualization for researchers, educators, and developers. Protein Science 30, 70–82 (2021).

64. Afonine, P. V. et al. Towards automated crystallographic structure refinement with *phenix.refine*. Acta Crystallogr D Biol Crystallogr 68, 352–367 (2012).

65. Emsley, P., Lohkamp, B., Scott, W. G. & Cowtan, K. Features and development of *Coot*. Acta Crystallogr D Biol Crystallogr 66, 486–501 (2010).

66. Varadi, M. et al. AlphaFold Protein Structure Database: massively expanding the structural coverage of protein-sequence space with high-accuracy models. Nucleic Acids Res 50, D439–D444 (2022).

67. Mirdita, M., et al. ColabFold - Making Protein Folding Accessible to All. 2021.08.15.456425 https://www.biorxiv.org/content/10.1101/2021.08.15.456425v2 (2021) doi:10.1101/2021.08.15.456425.

68. Jumper, J. et al. Highly accurate protein structure prediction with AlphaFold. Nature 596, 583–589 (2021).

69. Stariha, J. T. B., Hoffmann, R. M., Hamelin, D. J. & Burke, J. E. Probing Protein-Membrane Interactions and Dynamics Using Hydrogen-Deuterium Exchange Mass Spectrometry (HDX-MS). Methods Mol Biol 2263, 465–485 (2021).

70. Masson, G. R. et al. Recommendations for performing, interpreting and reporting hydrogen deuterium exchange mass spectrometry (HDX-MS) experiments. Nat. Methods 16, 595–602 (2019).

71. Perez-Riverol, Y. et al. The PRIDE database resources in 2022: a hub for mass spectrometry-based proteomics evidences. Nucleic Acids Res 50, D543–D552 (2022).

72. Bates, T. A. et al. Biolayer interferometry for measuring the kinetics of protein-protein interactions and nanobody binding. Nat Protoc 20, 861–883 (2025).

